# PQLC2 signals lysosomal cationic amino acid abundance to the C9orf72 complex

**DOI:** 10.1101/670034

**Authors:** Joseph Amick, Arun Kumar Tharkeshwar, Gabriel Talaia, Shawn M. Ferguson

**Affiliations:** Department of Cell Biology, Department of Neuroscience, Program in Cellular Neuroscience, Neurodegeneration and Repair, Yale University School of Medicine, New Haven, CT, 06510

## Abstract

The C9orf72 protein is required for normal lysosome function. In support of such functions, C9orf72 forms a heterotrimeric complex with SMCR8 and WDR41 that is recruited to lysosomes when amino acids are scarce. These properties raise questions about the identity of the lysosomal binding partner of the C9orf72 complex and the amino acid sensing mechanism that regulates C9orf72 complex abundance on lysosomes. We now demonstrate that an interaction with the lysosomal cationic amino acid transporter PQLC2 mediates C9orf72 complex recruitment to lysosomes. This is achieved through an interaction between PQLC2 and WDR41. The interaction between PQLC2 and the C9orf72 complex is negatively regulated by arginine, lysine and histidine, amino acids that PQLC2 transports across the membrane of lysosomes. These results define a new role for PQLC2 in the regulated recruitment of the C9orf72 complex to lysosomes and reveal a novel mechanism that allows cells to sense and respond to changes in the availability of cationic amino acids within lysosomes.

## Introduction

A hexanucleotide repeat expansion in a non-coding region of the C9orf72 gene causes familial forms of amyotrophic lateral sclerosis and frontotemporal dementia (DeJesus-Hernandez et al., 2011; Gijselinck et al., 2012; Renton et al., 2011). Although the repeat expansion results in a reduction in C9orf72 mRNA and protein levels, the extent to which this is relevant for disease pathogenesis remains unclear (Belzil et al., 2013; DeJesus-Hernandez et al., 2011; Gijselinck et al., 2012; Viode et al., 2018; Waite et al., 2014; Xi et al., 2013). Nonetheless, investigation of this topic has established that the C9orf72 protein is required for normal lysosome homeostasis in a variety of model systems including: mice, *C. elegans* and cultured human cells (Amick et al., 2016; Corrionero and Horvitz, 2018; McAlpine et al., 2018; O’Rourke et al., 2016; Sullivan et al., 2016; Zhang et al., 2018). Thus, beyond potential disease implications, C9orf72 has an evolutionarily conserved role in supporting lysosome function. The C9orf72 protein is unstable on its own and functions as part of a larger protein complex that also contains SMCR8 and WDR41 (Amick et al., 2016; Sellier et al., 2016; Sullivan et al., 2016; Ugolino et al., 2016; Xiao et al., 2016; Zhang et al., 2018). Consistent with a direct function at lysosomes, this C9orf72 protein complex localizes to the cytoplasmic surface of lysosomes (Amick et al., 2016; Amick et al., 2018). Interestingly, interactions between the C9orf72 complex and lysosomes are acutely regulated by changes in amino acid availability and are most prominent when cells are deprived of amino acids (Amick et al., 2016; Amick et al., 2018). Although there is now considerable evidence for a lysosomal site of action, the mechanisms that support the regulated recruitment of the C9orf72 complex to lysosomes in response to amino acid scarcity have remained unsolved.

In addition to their classically defined role in the break-down of cellular macromolecules, lysosomes also possess signaling functions. This is best appreciated in the context of the mTORC1 signaling pathway, a central pathway for the regulation of cell growth and metabolism (Shimobayashi and Hall, 2014). Amino acid availability is a key factor regulating mTORC1 activation. mTORC1 is recruited to lysosomes via binding to the Rag GTPases, whose nucleotide loading status is regulated by amino acid availability (Bar-Peled and Sabatini, 2014). Multiple lysosomal proteins have been identified that function as sensors for specific amino acids and which are integrated into signaling pathways that converge on the Rags (Wolfson and Sabatini, 2017). Although investigation of mTORC1 regulation has contributed significantly to our understanding of how amino acid availability is sensed at lysosomes and communicated to mTORC1 via the Rag GTPases, C9orf72 has not been implicated in the Rag GTPase pathway. This suggests the existence of an undefined amino acid-sensing mechanism that operates upstream of the C9orf72 complex at lysosomes.

In this study we identify proline glutamine loop containing 2 (PQLC2), a lysosomal transporter of cationic amino acids (Jezegou et al., 2012; Liu et al., 2012), as an interactor of the C9orf72 complex and find that PQLC2 mediates the recruitment of the C9orf72 complex to lysosomes. The interaction between PQLC2 and the C9orf72 complex is specifically regulated by availability of arginine, lysine and histidine – the amino acids transported by PQLC2. Collectively, these results define a mechanism for amino acid-regulated recruitment of the C9orf72 complex to lysosomes and reveal a new pathway whereby the status of the lysosomal lumen is communicated to the rest of the cell.

## Results

### Identification of PQLC2 as a lysosomal binding partner for the C9orf72 complex

To understand how the C9orf72 complex is recruited to lysosomes in response to amino acid starvation, we sought to identify lysosomal binding partners of the C9orf72 complex. Although the endogenous C9orf72 protein is found on lysosomes in starved cells (Amick et al., 2016; Amick et al., 2018), its low expression levels raised challenges for the detection of lysosomal interacting partners of C9orf72. We therefore took advantage of a lysosome-targeted C9orf72 chimera, which was made by fusing the first 39 amino acids of p18/LAMTOR1 to C9orf72-GFP (Fig. 1A). This region of LAMTOR1 contains an acidic dileucine motif, palmitoylation and myristoylation sites, and has been used previously to target proteins fused to it to the cytoplasmic surface of lysosomes (Amick et al., 2018; Menon et al., 2014; Nada et al., 2009). Fusion of this region of LAMTOR1 to C9orf72-GFP was previously demonstrated to support the efficient targeting of a functional C9orf72 protein to the cytoplasmic surface of lysosomes (Amick et al., 2018). We therefore used lysosome-targeted C9orf72 (Lyso-C9orf72-GFP) as bait in immunoprecipitation experiments to identify lysosome localized binding partners of the C9orf72 complex (Fig. 1B). Parallel anti-GFP IPs from cells co-expressing Lyso-C9orf72-GFP and SMCR8-tdTomato or control cells expressing lysosome targeted GFP (Lyso-GFP, Fig. 1A) and SMCR8-tdTomato were analyzed by mass spectrometry. This approach identified multiple proteins that were selectively enriched in the Lyso-C9orf72-GFP sample (Fig. 1C). This included known C9orf72 interactors such as: SMCR8, WDR41, and the ULK1 complex (ULK1, RB1CC1, ATG13 and ATG101) (Amick et al., 2016; Jung et al., 2017; Sellier et al., 2016; Sullivan et al., 2016; Xiao et al., 2016; Yang et al., 2016). We also identified PQ loop containing 2 (PQLC2), an amino acid transporter responsible for the efflux of cationic amino acids from lysosomes, in the Lyso-C9orf72-GFP samples (Jezegou et al., 2012; Liu et al., 2012) (Fig. 1C). This amino acid transporter was particularly interesting given that C9orf72 complex abundance at lysosomes is regulated by changes in amino acid availability (Amick et al., 2016; Amick et al., 2018).

**Figure 1.**
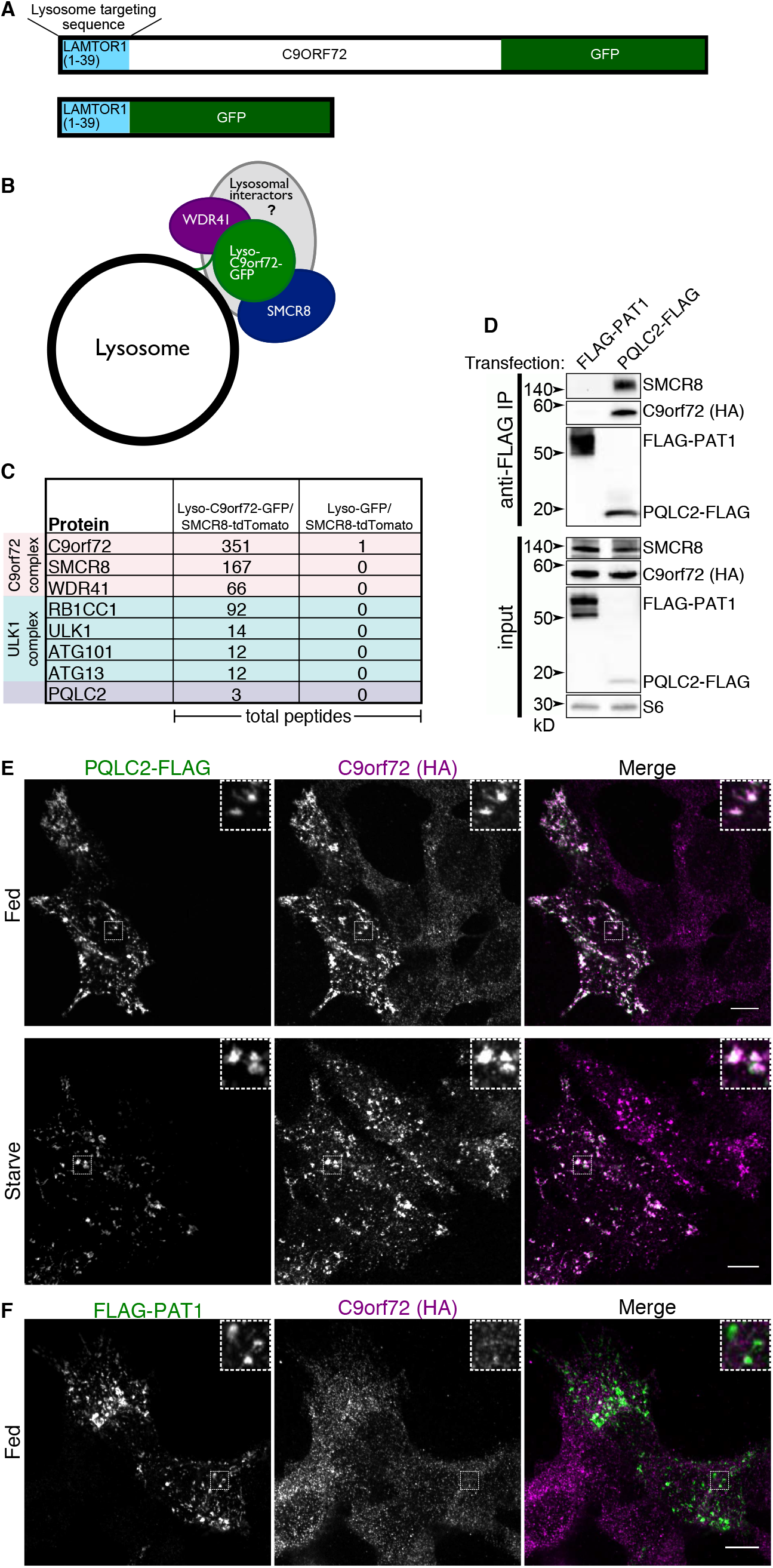
Identification of the amino acid transporter PQLC2 as a C9orf72 interacting protein. **(A)** Schematic diagram illustrating the lyso-C9orf72-GFP and the lyso-GFP construct used for proteomics. **(B)** Strategy to enrich the C9orf72 complex at lysosomes to attempt to identify lysosomal interacting partners. **(C)** Summary of proteins identified by LC-MS/MS that interact with lyso-C9orf72-GFP compared to a lyso-GFP control. Total peptide numbers are shown. **(D)** HEK293FT cells that express 2xHA-tagged C9orf72 from the endogenous locus were transfected with FLAG-tagged PQLC2 or lysosomal amino acid transporter PAT1, followed by anti-FLAG immunoprecipitation and immunoblotting for FLAG or endogenously expressed SMCR8 and C9orf72. **(E)** Effect of PQLC2 overexpression on C9orf72 localization in fed and starved cells. **(F)** Effect of PAT1 overexpression on C9orf72 localization in fed cells. Scale bars: 10 μm.

To further evaluate the interaction between PQLC2 and the C9orf72 complex, we performed immunoprecipitations on cells transfected with FLAG-tagged PQLC2 and found that it interacts with the endogenously-expressed C9orf72 and SMCR8 (Fig. 1D). As evidence of the specificity for the binding between PQLC2 and the C9orf72 complex, no interaction was detected between C9orf72 or SMCR8 and FLAG-tagged PAT1 (also known as LYAAT1 and SLC36A1), a lysosomal amino acid transporter of small neutral amino acids (Fig. 1D) (Sagne et al., 2001). Both the PQLC2-FLAG and FLAG-PAT1 proteins colocalize with LAMP1 on late endosomes and lysosomes, (Fig. S1A, B).

To assess colocalization between C9orf72 and PQLC2, cells that express 2xHA-tagged C9orf72 from the endogenous locus were transiently transfected to express FLAG-tagged PQLC2 under the control of the strong cytomegalovirus (CMV) promoter. In starved cells, C9orf72 localizes to PQLC2-positive lysosomes (Fig. 1E). Surprisingly the overexpression of PQLC2 also drove localization of C9orf72 to lysosomes even in fed cells (Fig. 1E). In contrast, overexpression of another lysosomal amino acid transporter, PAT1, did not noticeably alter C9orf72 localization (Fig. 1F). The interaction between C9orf72 and PQLC2 and the powerful effects of PQLC2 overexpression on C9orf72 localization suggested that interactions with PQLC2 control the recruitment of C9orf72 to lysosomes.

### PQLC2 is required for C9orf72 complex association with lysosomes

To test if PQLC2 is essential for C9orf72 recruitment to lysosomes, we used CRISPR-Cas9 to knock out PQLC2 in HEK293FT cells. After isolating clonal cell populations, we sequenced the PQLC2 locus and identified a line with frameshift deletions (7 bp and 34 bp deletions at the sgRNA target site in PQLC2 exon 2) and the absence of any remaining wildtype PQLC2 sequences. We then performed immunofluorescence analysis and found that C9orf72 was no longer recruited to lysosomes in response to starvation in PQLC2 knockout cells (Fig. 2A, B).

**Figure 2.**
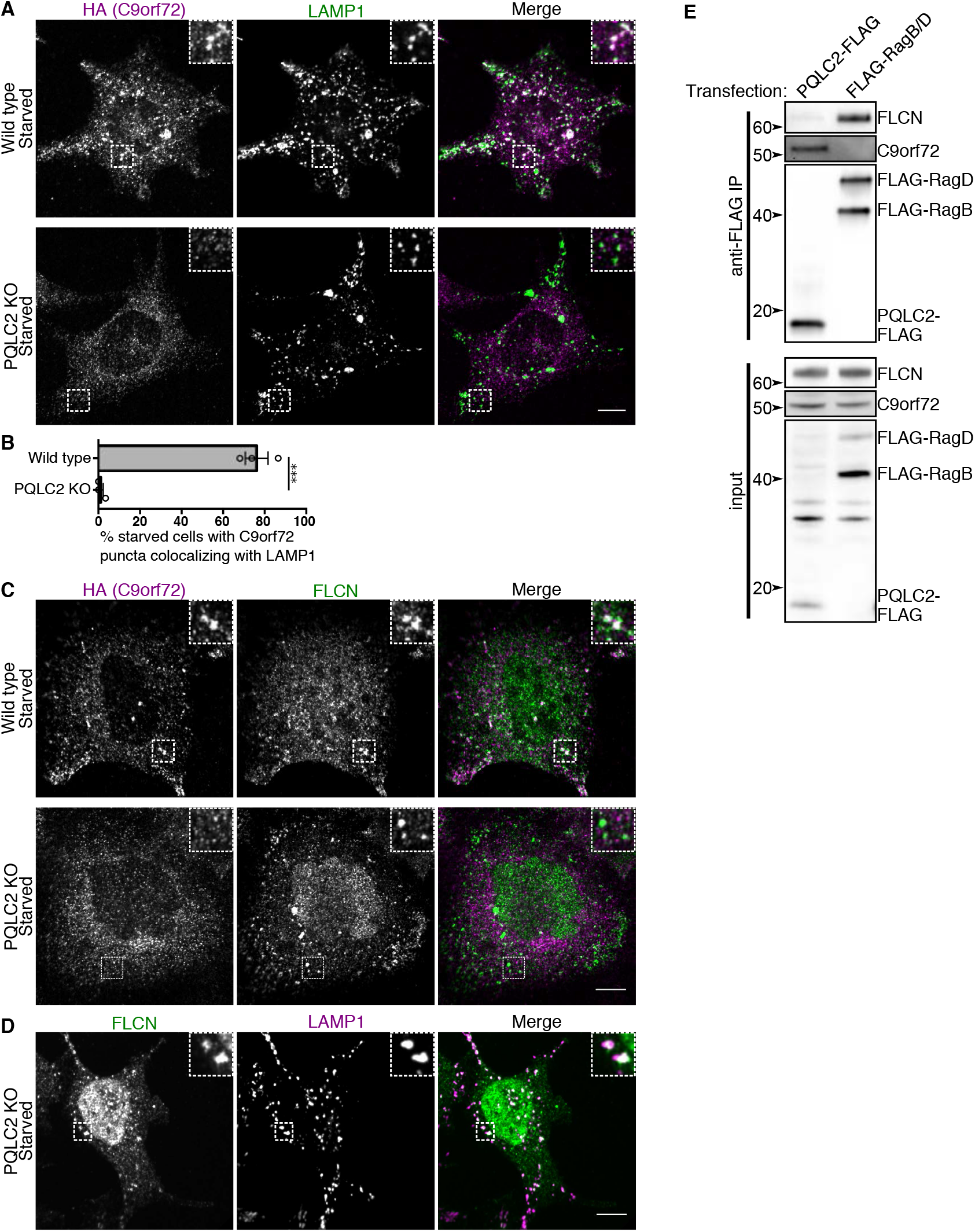
PQLC2 is required for C9orf72 recruitment to lysosomes. **(A)** Immunofluorescence images of C9orf72 localization (endogenously expressed 2xHA-C9orf72) in starved wild-type and PQLC2 knockout (KO) cells. Localization of C9orf72 to lysosomes (LAMP1) is lost in PQLC2 knockout cells. **(B)** For the indicated cell lines, the percentage of cells in starved conditions containing C9orf72 puncta that colocalize with LAMP1. Mean ± SEM is plotted with individual points displayed as open dots. Unpaired t test, p=0.0002. **(C)** Immunofluorescence images of C9orf72 and FLCN localization in starved wild type and PQLC2 knockout cells. **(D)** Immunofluorescence images of FLCN localization in starved PQLC2 knockout cells. Recruitment of FLCN to lysosomes (LAMP1 marker) is maintained in these cells. Scale bars: 10 μm. **(E)** Cells expressing PQLC2-FLAG and FLAG-tagged Rags B and D were subjected to anti-FLAG IPs and immunoblotting for FLAG and endogenous C9orf72 and FLCN.

C9orf72 and SMCR8 are predicted to be structurally similar to the folliculin (FLCN) and FLCN-interacting proteins (FNIPs) which also form a complex that is recruited to lysosomes in starved cells (Amick et al., 2016; Amick et al., 2018; Meng and Ferguson, 2018; Petit et al., 2013). To test the specificity of the requirement for PQLC2 in the lysosomal recruitment of the C9orf72 complex, we next examined FLCN localization in PQLC2 knockout cells using a FLCN antibody that was previously established to yield a specific lysosomal immunofluorescent signal (Meng and Ferguson, 2018). Although C9orf72 and FLCN both exhibit a punctate distribution in starved wild type cells, only FLCN still shows this punctate, LAMP1 co-localized, distribution in the PQLC2 knockouts (Fig 2C and D). These experiments reveal specificity in the role for PQLC2 on C9orf72 regulation.

Amino acid availability is communicated to FLCN via amino acid sensors upstream of the GATOR1 complex; cells lacking the Nprl3 subunit of GATOR1 are unable to recruit FLCN to lysosomes (Meng and Ferguson, 2018). To test the role for GATOR1 in communicating amino acid availability to C9orf72, we next knocked out Nprl3 in the background of a CRISPR knockin cell line that expresses 2xHA-C9orf72 from the endogenous locus. As expected, these cells are unable to efficiently inactivate mTORC1 during amino acid starvation (Fig S2A) (Bar-Peled et al., 2013; Panchaud et al., 2013). Unlike FLCN, C9orf72 was still recruited to lysosomes in starved Nprl3 KO cells (Fig S2B, C). Thus, while both FLCN and C9orf72 are recruited to lysosomes in response to amino acid starvation, they are recruited via different mechanisms. In addition, C9orf72 co-immunoprecipitates with PQLC2, while FLCN does not and FLCN co-immunoprecipitates with RagB and D, while C9orf72 does not (Fig 2E). These results are consistent with a PQLC2-dependent lysosome recruitment mechanism for C9orf72 that is distinct from the Rag-dependent recruitment mechanism for FLCN.

### Conserved prolines in PQLC2 are required for its interaction with the C9orf72 complex

PQLC2 contains two conserved PQ motifs (Fig S3A). In related transporters, the proline residues within this motif act as hinges within transmembrane helices that support conformational changes that are critical for the transport cycle (Lee et al., 2015). Replacement of these prolines with leucines was previously shown to nearly abolish PQLC2 transport of amino acids (Liu et al., 2012). We therefore tested if these residues are required for the interaction between PQLC2 and the C9orf72 complex. After confirming that the PQLC2 P55L, P201L and P55L/P201L double mutants localize to lysosomes (Fig S3B-D), we next performed immunoprecipitations to test their ability to interact with the C9orf72 complex. C9orf72, SMCR8 and WDR41 co-purified with wild type PQLC2 to a similar degree in fed and starved conditions, while the interactions between the C9orf72 complex and the PQLC2 P55L, P201L and P55L/P201L double mutants were greatly reduced (Fig. 3A).

**Figure 3.**
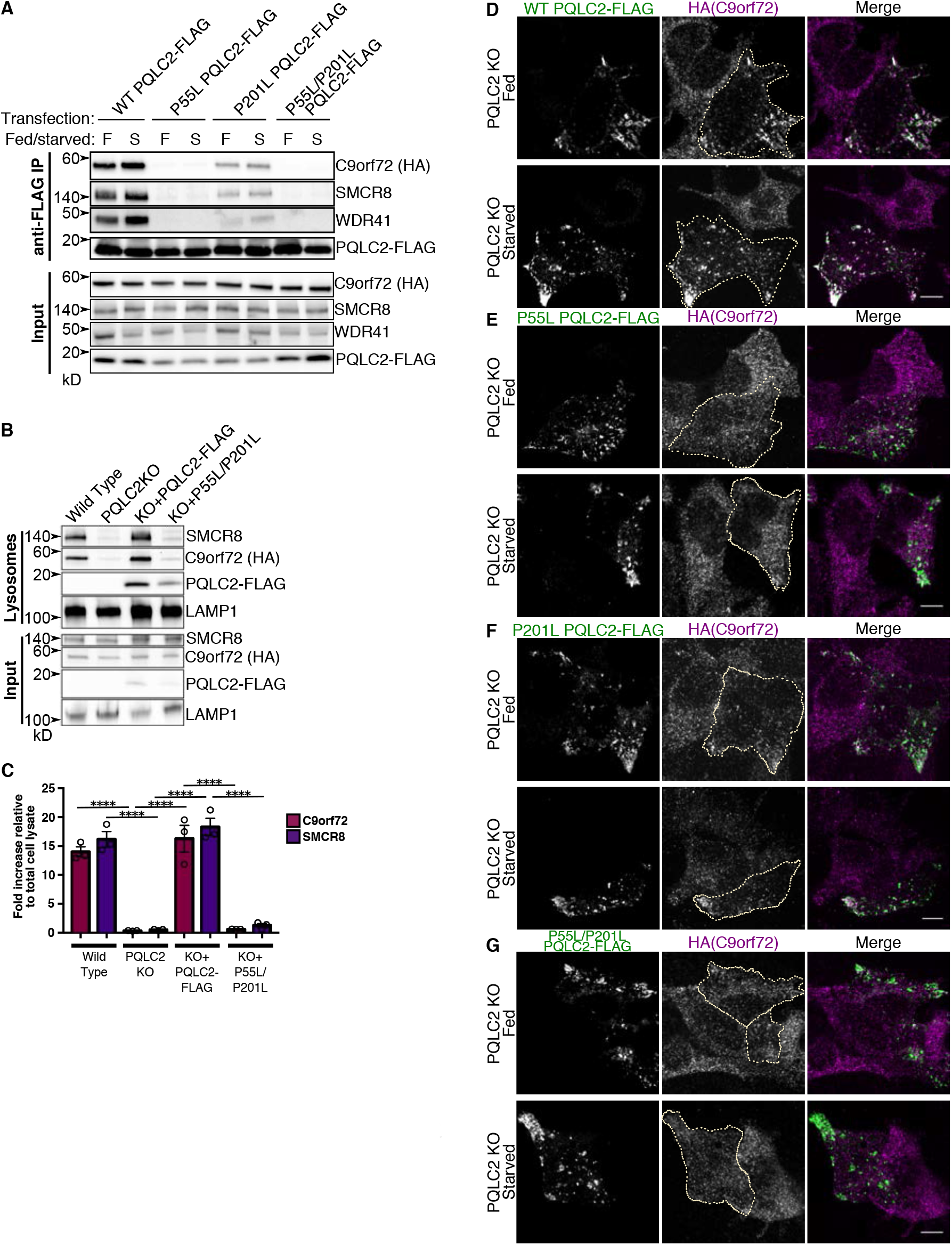
PQ motif mutations in PQLC2 disrupt C9orf72 complex interactions and recruitment of C9orf72 to lysosomes. **(A)** PQLC2 knockout cells were transfected with wild type (WT) PQLC2-FLAG or the indicated mutants. Cells were cultured under fed (F) or starved (S) conditions; PQLC2-FLAG was immunoprecipitated, followed by immunoblotting for FLAG and endogenous WDR41, C9orf72 and SMCR8. **(B)** Lysosomes were purified from the parental HEK293FT cells, PQLC2 knockout cells, and PQLC2 knockout cells stably expressing wild type PQLC2-FLAG or P55L/P201L PQLC2-FLAG. Immunoblots of the indicated proteins in the total cell lysate (input) and in magnetically isolated lysosomes (lysosomes). **(C)** Quantification of immunoblots for the indicated proteins expressed as a fold increase in the lysosome fraction relative to the input (mean ± SEM, three independent experiments, individual data points shown as open circles; ****P ≤ 0.0001, ANOVA with Tukey’s multiple comparisons test). **(D-G)** PQLC2 knockout cells were transfected with wild type (WT) PQLC2-FLAG, or the indicated mutants and cultured in normal growth medium (fed) or amino acid/serum-free RPMI (starved). Cells were fixed and immunostained with FLAG and HA antibodies to examine C9orf72 localization. Transfected cells are outlined. Scale bars: 10 μm.

As a complementary approach to test the role of PQLC2 and PQ loop mutants in the recruitment of C9orf72 to lysosomes, we analyzed lysosomes purified from wild type, PQLC2 knockout, and knockout cells reconstituted with wild type PQLC2-FLAG or the P55L/P20L double mutant. Lysosomes were endocytically loaded with iron dextran nanoparticles and then subjected to a magnetic purification protocol (Amick et al., 2018; Tharkeshwar et al., 2017). Following lysosome isolation, C9orf72 and SMCR8 were detected on lysosomes in wild type, but not PQLC2 KO cells (Fig. 3B, C). C9orf72 and SMCR8’s presence on lysosomes was restored when wild type PQLC2-FLAG was added back to knockout cells. However, the P55L/P201L mutant failed to rescue recruitment of the C9orf72 complex to lysosomes (Fig. 3B, C). The impact of PQLC2 PQ-motif mutations on lysosome recruitment of C9orf72 was further corroborated by immunofluorescence experiments. Although wild type PQLC2 conferred constitutive lysosome localization to C9orf72 in the PQLC2 knockout cells, expression of P55L or P201L single mutants in PQLC2 knockout cells showed a much-reduced ability to support C9orf72 recruitment to lysosomes in either the fed or starved states (Fig 3D-F) while the P55L/P201L double mutant failed to recruit C9orf72 (Fig 3G). Collectively, the results of immunofluorescence, lysosome purification and co-IP experiments establish that: 1) The interaction between PQLC2 and the C9orf72 complex is essential for its recruitment to lysosomes; and 2) the PQ motifs that are predicted to support PQLC2 conformational changes that accompany substrate transport are also critical for supporting C9orf72 complex binding to PQLC2.

### Cationic amino acids regulate the interaction between PQLC2 and the C9orf72 complex

Localization of the C9orf72 complex to lysosomes is regulated by amino acid availability; this implies that the interaction with PQLC2 should also be regulated by amino acid availability. However, overexpressed PQLC2 interacted with the C9orf72 complex constitutively, with only a modest difference between fed and starved conditions (Fig. 1E, 3A, 3D). To overcome issues related to over-expression, we used genome editing to insert a 2xHA epitope tag into the endogenous PQLC2 gene in HEK293FT cells to yield cells that express PQLC2 with a C-terminal 2xHA epitope tag from the endogenous locus. The successful insertion of the tag was confirmed by sequencing of genomic DNA (Fig S4A-B). The PQLC2-2xHA cell line yielded a specific HA immunofluorescent signal compared to the parental cell line (Fig S4C). As expected, PQLC2-2xHA localized robustly to lysosomes (Fig S4D). We then immunoprecipitated PQLC2 from these cells under different nutrient conditions. Cells were cultured in media with or without dialyzed serum and/or the minimal essential medium (MEM) amino acids mix. The withdrawal of amino acids, but not serum, stimulated the interaction between PQLC2 and all three components of the C9orf72 complex (Fig 4A, B). We next asked which amino acids regulate the interaction. The MEM mix contains 12 amino acids, including the substrates for PQLC2 (arginine, lysine and histidine; Fig 4C). Cells were cultured in media containing the full amino acid mix, media lacking all amino acids, media lacking the 9 non-arginine/lysine/histidine amino acids in the full mix, or media lacking just arginine, lysine and histidine. Relative to cells cultured in the complete MEM amino acids media, the interaction with PQLC2 was increased in cells starved of all amino acids or starved of just arginine, lysine and histidine. However, the interactions was not stimulated when cells were starved of the other 9 MEM amino acids (Fig 4D, E).

**Figure 4.**
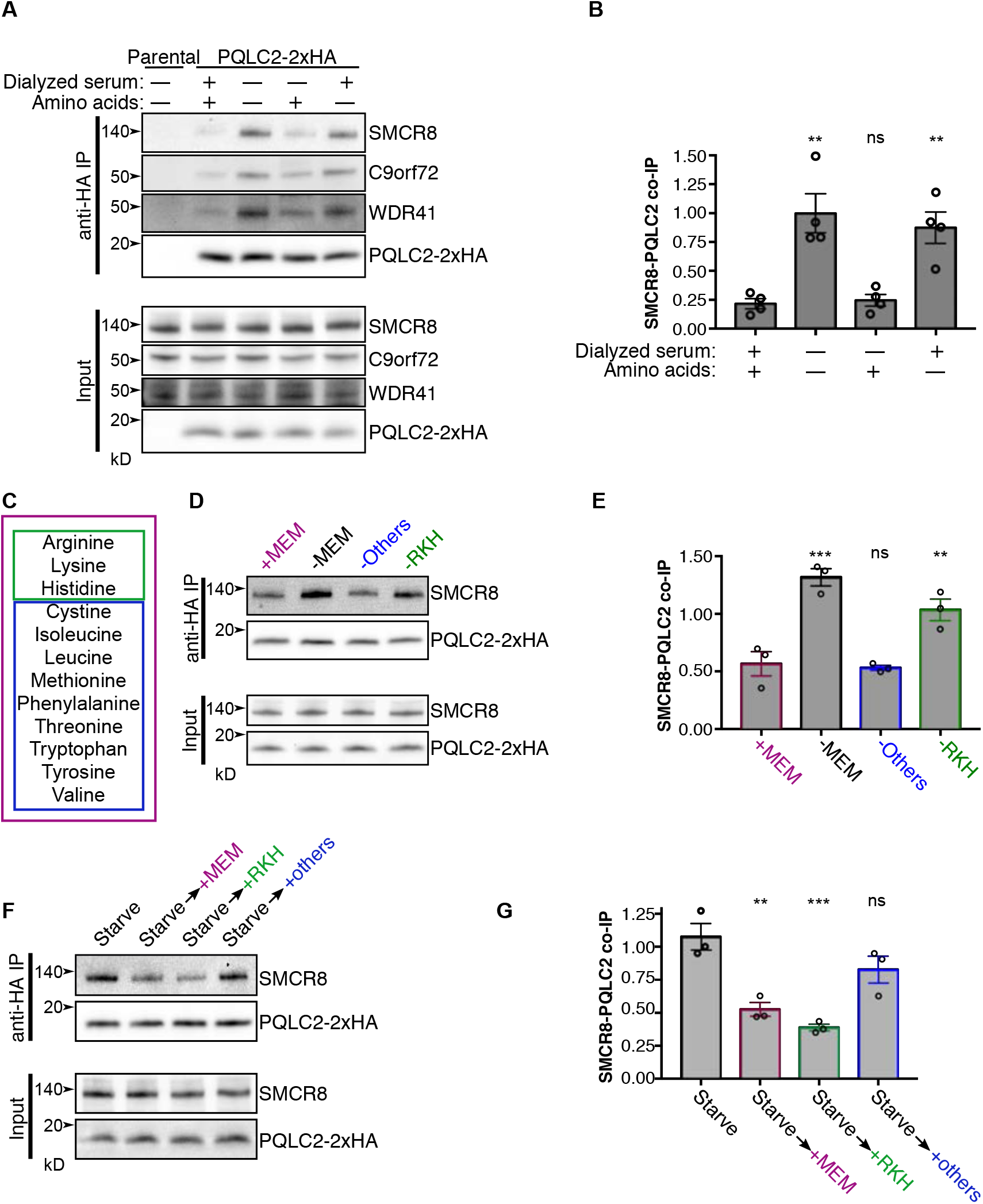
Amino acid dependent regulation of the C9orf72 complex’s interaction with PQLC2. **(A)** Wild type (parental) or gene-edited cells expressing PQLC2-2xHA from the endogenous locus were incubated in medium with or without amino acids and/or dialyzed serum. PQLC2 was then immunoprecipitated followed by immunoblotting with the indicated antibodies. **(B)** Summary of the ratio of SMCR8 to PQLC2-2xHA in IPs in **(A)**. All values are mean ±SEM with individual data points indicated by open circles. ** P ≤ 0.01, ns, not significant (ANOVA with Dunnett’s multiple comparisons test). **(C)** List of amino acids in minimal essential medium amino acid mix. Magenta, green and blue boxes indicate groups of amino acids used in subsequent experiments. **(D)** Cells incubated in media containing the MEM amino acid mix (+MEM, magenta box), media without amino acids (-MEM), without arginine, lysine and histidine (-RKH, green box), or without the other amino acids in the MEM mix (-others, blue box). PQLC2 was then immunoprecipitated to determine the effects of these conditions on the interaction. **(E)** Summary of the ratio of SMCR8 to PQLC2-2xHA in IPs in **(D)**. ** P ≤ 0.01, *** P ≤ 0.001, ns, not significant (ANOVA with Dunnett’s multiple comparisons test). **(F)** Cells were starved, then incubated in media containing the indicated amino acids, followed by immunoprecipitation of PQLC2. **(G)** Summary of the ratio of SMCR8 to PQLC2-2xHA in IPs in **(F)**. ** P ≤ 0.01, *** P ≤ 0.001, ns, not significant (ANOVA with Dunnett’s multiple comparisons test).

We then performed the reciprocal experiment wherein cells were first starved of all amino acids to promote interactions between PQLC2 and the C9orf72 complex and then measured the response to the re-feeding of specific amino acids. The complete MEM mixture reduced the co-IP between the C9orf72 complex and PQLC2 as did the mix of arginine, lysine and histidine (Fig 4F, G). In contrast, addition of the nine remaining amino acids did not significantly alter this interaction. Therefore, cationic amino acid (arginine, lysine and histidine) availability specifically regulates the interaction between PQLC2 and components of the C9orf72 complex. Consistent with the analysis of interactions between the C9orf72 complex and PQLC2, immunofluorescence analysis revealed that C9orf72 recruitment to lysosomes was reduced following addition to starved cells of the complete MEM amino acid mix and by a mix of arginine, lysine and histidine, but not by the other amino acids in the MEM mixture (Fig. 5). These results suggest that the availability of its substrate directly regulates the ability of the PQLC2 to recruit the C9orf72 complex to lysosomes. An alternative possibility is that substrate availability affects the lysosomal localization of PQLC2 itself. However, this was examined and we did not observe any impact of amino acid starvation or feeding on the lysosomal enrichment of PQLC2 (Fig S5).

**Figure 5.**
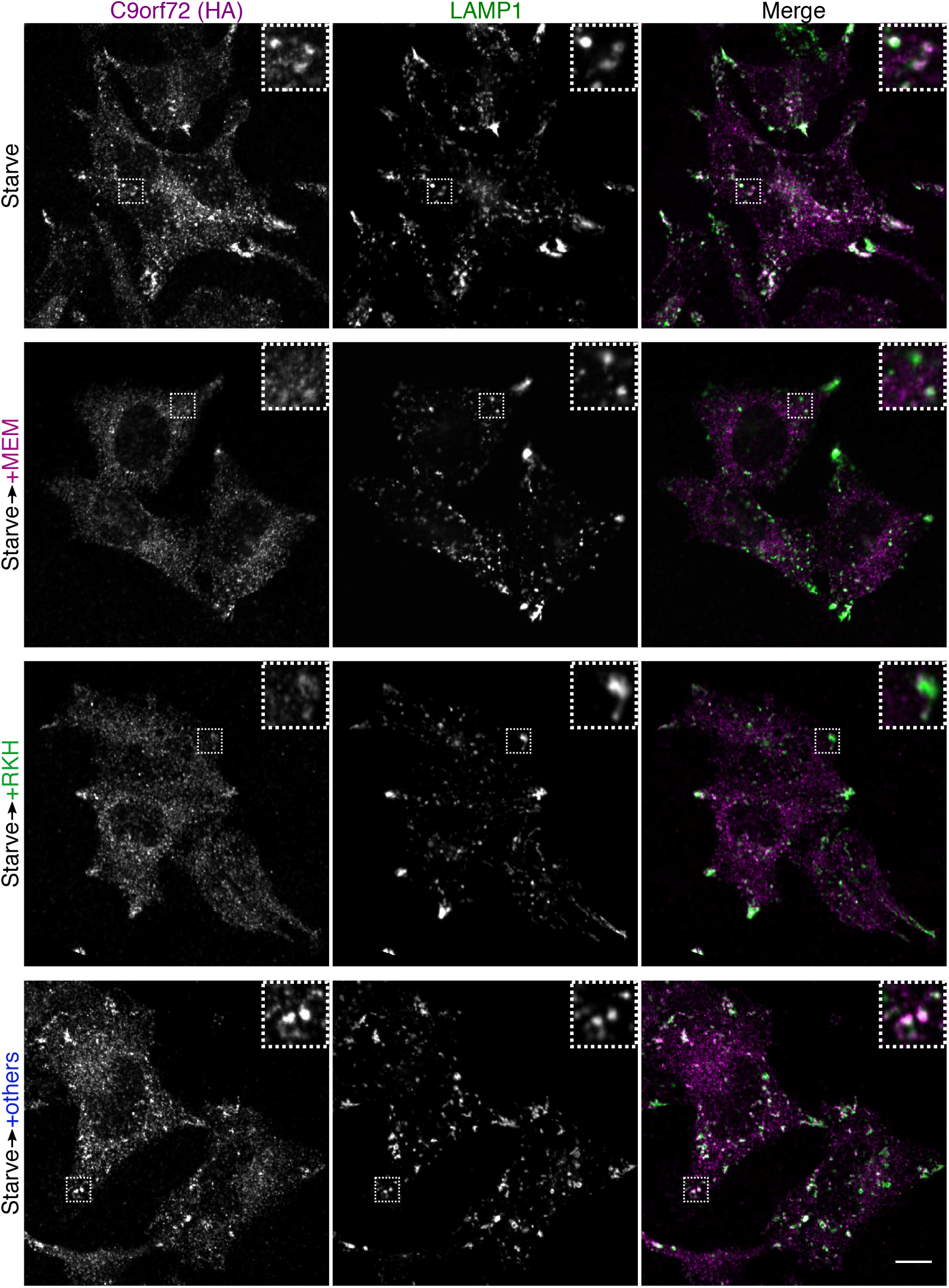
Arginine, lysine and histidine regulate C9orf72 localization. Immunofluorescence images of 2xHA-C9orf72 and LAMP1 in cells expressing 2xHA-C9orf72 from the endogenous locus. Cells were starved, then incubated in media containing the indicated amino acids (as in Figure 5 C, F, G). Scale bar: 10 μm.

The observation that the interaction between the C9orf72 complex and PQLC2 is regulated by availability of arginine, lysine and histidine, the amino acids transported out of lysosomes by PQLC2, suggested that PQLC2 communicates luminal amino levels to the C9orf72 complex. To more directly test this model, we designed and implemented a strategy to acutely generate a PQLC2 substrate in the lumen of lysosomes. This took advantage of a known relationship between PQLC2 and cystinosin, the lysosomal transporter for cystine (Kalatzis et al., 2001). Mutations in cystinosin cause cystinosis, a lysosome storage disorder characterized by the massive accumulation of cystine within lysosomes (Gahl, 2009; Gahl et al., 1982; Town et al., 1998). Cysteamine is used to treat this disease (Markello et al., 1993; Thoene et al., 1976). It reacts with cystine in lysosomes by sulfhydryl-disulfide exchange to form cysteine and a mixed disulfide (Gahl, 2009; Thoene et al., 1976). The mixed disulfide is structurally similar to lysine, and is transported out of lysosomes by PQLC2 (Jezegou et al., 2012; Liu et al., 2012; Pisoni et al., 1985). Thus, adding cysteamine to cystinosin-depleted cells acutely generates a PQLC2 substrate within the lumen of lysosomes (Fig 6A).

**Figure 6.**
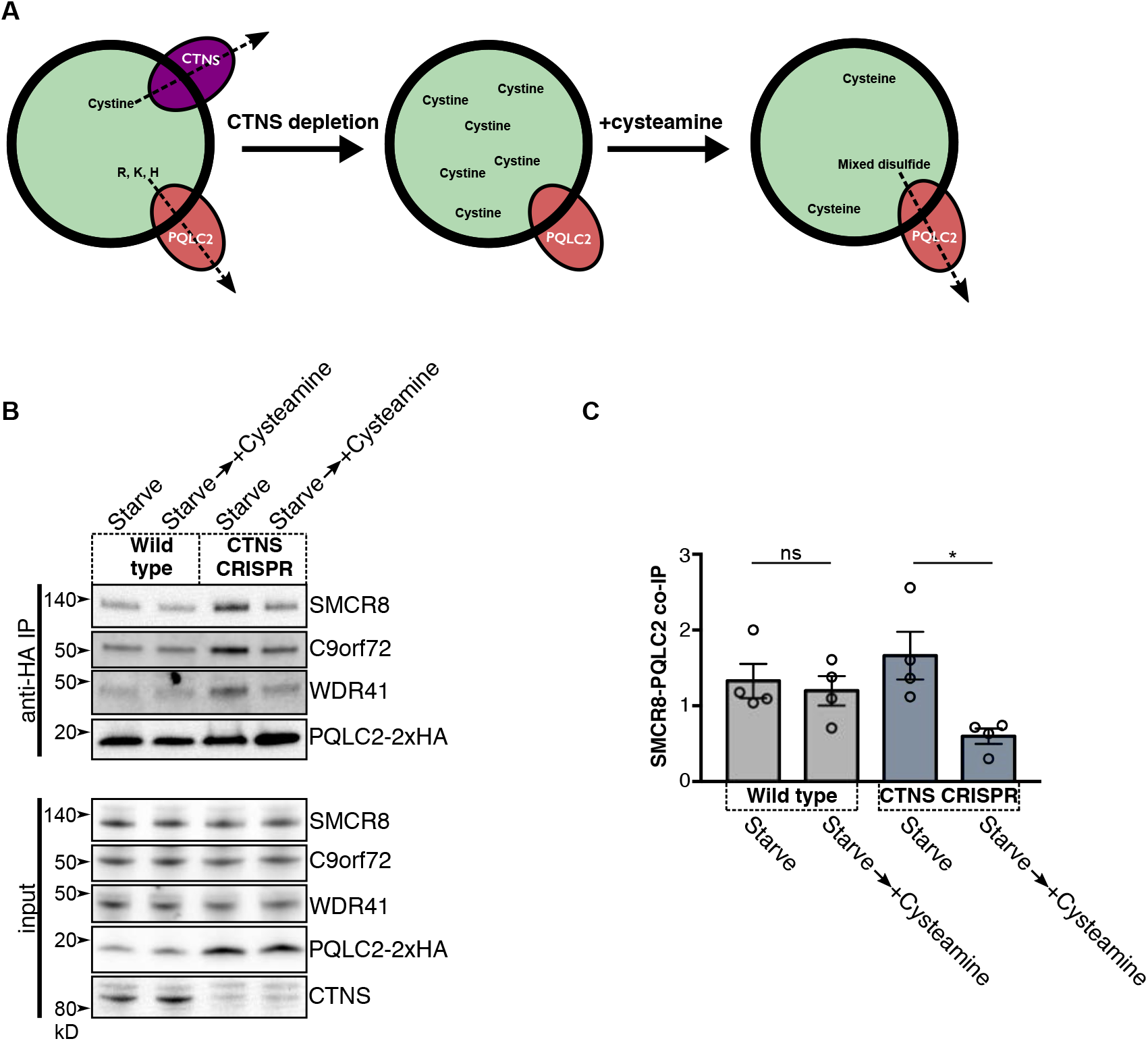
Lysosomal substrates negatively regulate PQLC2’s interaction with the C9orf72 complex. **(A)** Principle of the cystinosin depletion/cysteamine treatment experiment. Cystinosin (CTNS) is a lysosomal transporter of cystine and PQLC2 is a lysosomal transporter of arginine, lysine and histidine. Upon depletion of CTNS, cystine accumulates within lysosomes. When the drug cysteamine is added, it reacts with cystine in lysosomes to form cysteine and a mixed disulfide. The mixed disulfide is transported by PQLC2. Therefore, upon CTNS depletion, PQLC2 substrate can be generated within lysosomes by cysteamine addition and the effect on the interaction between PQLC2 and the C9orf72 complex assayed. **(B)** Wild type or CTNS-depleted cells expressing PQLC2-2xHA from the endogenous locus were starved, then incubated in starvation media containing cysteamine (60 minutes, 1 mM). PQLC2 was immunoprecipitated from cell lysates, followed by immunoblotting with the indicated antibodies. **(C)** Summary of the ratio of SMCR8 to PQLC2-2xHA in IPs in **(B)**. Values are mean ±SEM from four independent experiments with individual data points indicated by open circles. * P ≤ 0.05 (ANOVA with Tukey’s multiple comparisons test).

When wild type cells expressing PQLC2-2xHA from the endogenous locus were starved and treated with cysteamine the interaction between PQLC2 and the C9orf72 complex remained steady. This was expected given that cystinosin prevents the buildup of lysosomal cystine in these cells. However, in a genome edited cell line that was depleted of cystinosin, acute cysteamine treatment reversed the starvation-induced interaction between PQLC2 and the C9orf72 complex (Fig 6B, C). This result supports a model wherein PQLC2 communicates the abundance of its substrates within the lysosome lumen to the cytoplasmic C9orf72 complex.

### WDR41 mediates interaction between the C9orf72 complex and PQLC2

WDR41 was previously defined as required for the recruitment of C9orf72 and SMCR8 to lysosomes (Amick et al., 2018). We therefore hypothesized that WDR41 would be required for the interaction of the C9orf72 complex with PQLC2. This was tested in wild type versus WDR41 knockout cells (Amick et al., 2018) that were transfected with PQLC2-FLAG. PQLC2-FLAG coimmunoprecipitated endogenous WDR41, SMCR8 and C9orf72 in the wild type cells and these interactions between C9orf72 and SMCR8 and PQLC2 were abolished in WDR41 KO cells (Fig 7A, B). Imaging WDR41 knockout cells transfected with PQLC2-FLAG confirmed that PQLC2 still localized robustly to lysosomes in the absence of WDR41 (Fig 7C). Wild type and WDR41 KO cells were next transfected with PQLC2-FLAG and endogenous C9orf72 localization was examined by immunofluorescence. While C9orf72 localizes to PQLC2-positive lysosomes in wild type cells, this pattern of colocalization was absent in WDR41 knockouts (Fig 7D). Lastly, we asked if WDR41 itself was able to interact with PQLC2 in the absence of C9orf72 and SMCR8. To this end, wild type and C9orf72+SMCR8 double knockout cells (Amick et al., 2016) were transfected with PQLC2-FLAG. Anti-FLAG immunoprecipitations subsequently revealed that WDR41 was still able to interact with PQLC2 in the absence of both C9orf72 and SMCR8 (Fig 7E, F). However, the moderate reduction of this interaction in the double KO cell line indicates that C9orf72 and SMCR8 positively contribute to this binding. In summary, multiple pieces of evidence indicate that WDR41 is critical for the interaction between PQLC2 and the C9orf72 complex at lysosomes.

**Figure 7.**
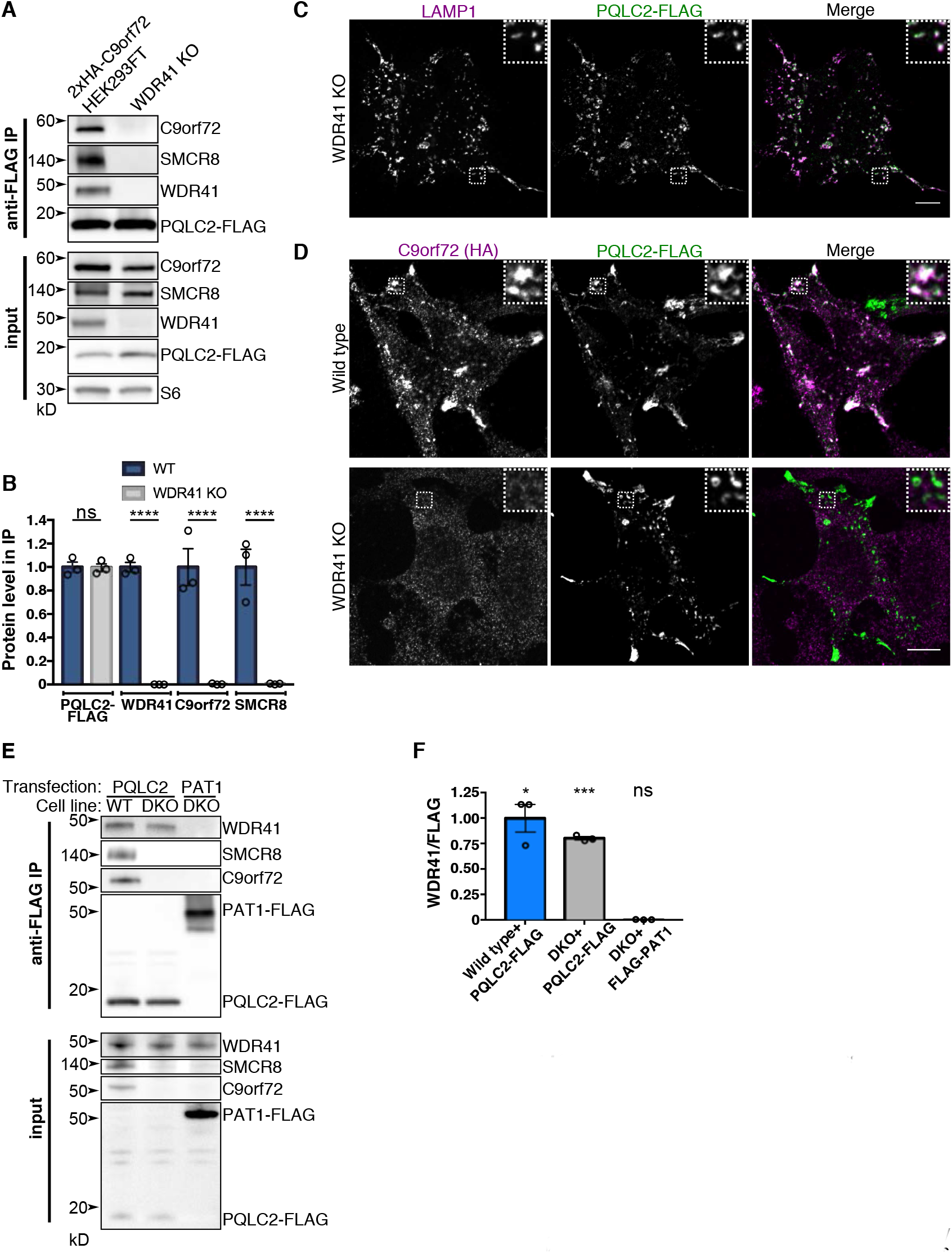
WDR41 mediates the interaction of the C9orf72 complex with PQLC2. **(A)** PQLC2-FLAG was expressed in wild type and WDR41 knockout cells. Following anti-FLAG immunoprecipitation, PQLC2-FLAG, and endogenous WDR41, C9orf72 and SMCR8 were detected by immunoblotting. **(B)** Summary of results from **(A)**. Level of the indicated proteins in anti-FLAG immunoprecipitations from wild type and WDR41 knockout cells (wild type levels normalized to 1, n=3, mean±SEM plotted with individual points as open circles). **(C)** Immunofluorescence of PQLC2-FLAG and LAMP1 in WDR41 knockout cells. **(D)** Immunofluorescence images of C9orf72 localization (endogenously expressed 2xHA-C9orf72) in wild type and WDR41 knockout cells transiently overexpressing PQLC2-FLAG. Consistent with IP data, overexpression of PQLC2 results in co-localization with C9orf72 in wild type but not WDR41 knockouts. **(E)** Wild type (WT) and C9orf72/SMCR8 double knockout cells (DKO) were transfected with FLAG-tagged PQLC2 and immunopreciptated. FLAG-tagged PAT1, a lysosomal amino acid transporter, served as a negative control. **(F)** Summary of WDR41 levels in anti-FLAG immunoprecipitations from **(E)**. n=3, mean ± SEM plotted with individual points as open circles. One-sample t test comparing to a hypothetical mean of 0; ns=not significant, * P ≤ 0.05, *** P ≤ 0.001.

## Discussion

It was previously established that C9orf72 is required for the maintenance of normal lysosome function and is itself part of a protein complex that localizes to the cytoplasmic surface of lysosomes when cells are starved of amino acids (Amick et al., 2016; Corrionero and Horvitz, 2018; McAlpine et al., 2018; O’Rourke et al., 2016; Sullivan et al., 2016; Zhang et al., 2018). These observations define two major questions concerning the pathway in which C9orf72 functions. First, what is the basis for the regulated interactions that connect the C9orf72 complex to lysosomes? Second, what is the specific function of this complex at lysosomes? In this study we solved the first of these two problems by identifying a regulated physical interaction between PQLC2, a lysosomal amino acid transporter, and the C9orf72 complex. We furthermore established that PQLC2 is both necessary and sufficient for recruiting the C9orf72 complex to lysosomes. This process is negatively regulated by the cationic amino acids that are transported by PQLC2. We thus identify a new mechanism whereby cationic amino acid availability is sensed at lysosomes. Importantly, this process is distinct from previously defined lysosome-based amino acid sensing mechanisms that converge on the Rag GTPases for the regulation of mTORC1 signaling.

Our results reveal that in addition to acting as a cationic amino acid transporter, PQLC2 also controls the abundance of the C9orf72 complex at lysosomes in response to changes in cationic amino acid availability. Other transporters with dual functionality related to substrate transport and signaling haven been termed “transceptors” (Holsbeeks et al., 2004). One example of a transceptor is the yeast general amino acid permease GAP1, which transports a broad range of amino acids and acts as an amino acid sensor that activates protein kinase A signaling (Donaton et al., 2003; Grenson et al., 1970). Meanwhile, at the mammalian lysosome, SLC38A9 is an amino acid transporter that associates with mTORC1 regulatory machinery and communicates the abundance of arginine, lysine and leucine to this signaling pathway (Jung et al., 2015; Rebsamen et al., 2015; Wang et al., 2015). Our new data supports the inclusion of PQLC2 into the group of transporters that have the ability to perform a second function.

PQLC2 is part of a family of transporters with distant homology to the bacterial rhodopsin family of receptors, which includes the SWEET/semiSweet family of sugar transporters (Xuan et al., 2013; Zhai et al., 2001). Crystal structures of *E. coli* semiSWEET at different steps of its transport cycle revealed that the conserved PQ motif acts as a hinge that allows the transporter to transition between different steps of its transport cycle (Lee et al., 2015). For transceptors, conformational changes that arise during the transport cycle are thought to be transduced to downstream effector proteins (Thevelein and Voordeckers, 2009). The interaction between PQLC2 and the C9orf72 complex is mediated by WDR41 and requires conserved proline residues within the PQ motifs of PQLC2. This suggests that the interaction between WDR41 and PQLC2 requires conformational changes that are enabled by these hinges in PQLC2. However, the identity of specific cytoplasmic residues on PQLC2 that might be revealed by such conformational changes have not yet been established. An additional possibility for how substrate availability could affect the ability of PQLC2 to interact with WDR41 could be via changes in PQLC2 oligomerization state. Such regulation would parallel reports of the regulation of presynaptic dopamine transporter oligomerization by its substrates (Chen and Reith, 2008). Further progress in this area will require defining the relevant binding sites on PQLC2 and WDR41, insight into structural changes arising in PQLC2 during its transport cycle, and how the presence or absence of amino acid substrates in the lysosome or cytosol alter these states.

We used cystinosin depletion combined with cysteamine treatment as a tool to acutely generate a PQLC2 substrate within the lysosome lumen as a strategy to test the effect of lysosomal substrate on the interaction between PQLC2 and the C9orf72 complex. The observation that this treatment negatively regulates the PQLC2-C9orf72 complex interaction also reveals a potential unanticipated effect of cysteamine treatment, which could merit investigation in the context of cystinosis disease therapy (Markello et al., 1993).

Like the C9orf72 complex, the FLCN-FNIP heterodimer also preferentially associates with lysosomes in response to amino acid depletion (Meng and Ferguson, 2018; Petit et al., 2013; Tsun et al., 2013). The FLCN-FNIP and the C9orf72 complexes both contain two interacting DENN domain proteins (Amick and Ferguson, 2017). However, these two complexes possess distinct mechanisms underlying their regulation. While FLCN-FNIP is recruited to lysosomes via a direct interaction with Rag GTPases (Meng and Ferguson, 2018; Petit et al., 2013; Tsun et al., 2013), the C9orf72 complex instead relies on the interaction between WDR41 and PQLC2. The interaction between FLCN-FNIP and the Rags is selective for the GDP-bound state of RagA/B and is thus dependent on GATOR1, the GAP that stimulates RagA/B GTPase activity (Bar-Peled et al., 2013). The specificity of these unique mechanisms for responding to changes in amino acid availability is illustrated by our observations that FLCN is still enriched on lysosomes following the starvation of PQLC2 KO cells (Fig. 2D) and the intact recruitment of C9orf72 to lysosomes in NPRL3 (GATOR1 subunit) KO cells (Fig. S2C). These results demonstrate that interactions between the C9orf72 complex and PQLC2 define a distinct and novel amino acid sensing pathway at the lysosome.

Both the C9orf72 and SMCR8 proteins are predicted to contain DENN domains (Levine et al., 2013; Zhang et al., 2012). In other proteins, DENN domains are best characterized for their role as guanine nucleotide exchange factors (GEFs) for small GTPases of the Rab family (Marat et al., 2011; Wu et al., 2011). However, although several small GTPases have been associated with the C9orf72 complex, including Rab1 (Webster et al., 2016), Rab3 (Frick et al., 2018), Rab5 (Shi et al., 2018), Rabs 8 and 39 (Sellier et al., 2016) and Arf6 (Sivadasan et al., 2016), none of them are known to have lysosome-localized functions that are regulated by either C9orf72 or SMCR8. Our new identification of a PQLC2-dependent mechanism that explains how the C9orf72 complex is recruited to lysosomes provides a foundation for the future efforts to identify the down-stream target(s) of the C9orf72 complex at lysosomes and to integrate this pathway into a broader understanding of how nutrient sensing at lysosomes is integrated into the control of cellular physiology.

In summary, we have identified the cationic amino acid transporter PQLC2 as a lysosome-localized binding partner for the C9orf72 complex that is essential for the recruitment of C9orf72, SMCR8 and WDR41 to lysosomes when cells are deprived of cationic amino acids. Our data are consistent with a model wherein conformational changes induced by the availability of luminal substrates are critical for the regulated interactions between PQLC2 and the C9orf72 complex. Collectively, these findings reveal a new amino-acid sensing mechanism at the lysosome.

## Acknowledgments

We appreciate lab management contributions of Agnes Ferguson that supported this project and are grateful to all members of the Ferguson lab for helpful advice and discussion. We thank Jean Kanyo of the Mass Spectrometry and Proteomics Resource of the W.M. Keck Foundation Biotechnology Resource Laboratory at Yale University for her work on mass spectrometry experiments. Mass spectrometry experiments were supported by NIH shared instrument grant 1S10OD018034-01. This research was supported by NIH grant GM105718 to SMF. JA was supported by grants T32GM007223 and F31GM119249 from the NIH.

## Methods

### Cell Culture and Transfection

HEK293FT cells (Life Technologies) were grown in Dulbecco’s Modified Eagle Medium (DMEM) +4.5 g/L D-glucose, L-Glutamine (Gibco), 10% fetal bovine serum, and 1% penicillin/streptomycin supplement (Mediatech). Transfections were performed with 4 μg of DNA, 800 μL of OptiMEM (Invitrogen, and 12 μL of FuGENE 6 transfection reagent (Promega) per 10 cm dish. Cells were analyzed two days post-transfection.

### Plasmids

PQLC2-FLAG plasmids were generated as follows. The PQLC2 coding sequence was purchased as a gBlock (Integrated DNA Technologies). The gBlock sequence is listed in Supplemental Table 1. This DNA was inserted into SmaI-digested pEGFPN2 vector (Clontech) by Gibson Assembly (NEBuilder HiFi DNA Assembly, New England BioLabs) according to manufacturer protocols. A FLAG-tag (DYKDDDDK tag) followed by a stop codon was then inserted using the Q5 Site-Directed Mutagenesis Kit (New England BioLabs) according to manufacturer protocols. Primers for this reaction are listed in Supplemental Table 2. PQLC2 mutants were also generated using the Q5 Site-Directed Mutagenesis Kit and primers for generating these mutants are listed in Supplemental Table 2. For lentivirus-mediated transgenic expression of PQLC2-FLAG and mutants, these sequences were amplified by PCR from plasmids described above and inserted into SmaI-digested pLVX-puro vector (Clontech) by Gibson Assembly. Sequences for these PCR primers are listed in Supplemental Table 2. Generation of the Lyso-C9orf72-GFP plasmid and the SMCR8-tdTomato plasmid was described previously (Amick et al., 2016; Amick et al., 2018). Stbl3 *E. coli* (Invitrogen) were used for cloning pLVX-puro plasmids, XL1-Blue E. coli (Agilent) were used for cloning other plasmids. All plasmids were sequence verified. Flag pLJM1 RagB and RagD plasmids were gifts from David Sabatini (Addgene plasmid 19313 and 19316) (Sancak et al., 2008).

### Immunoprecipitation and immunoblotting

Prior to plating cells, 10 cm dishes were coated with 2 μg/mL Poly-D lysine (Sigma-Aldrich) in PBS overnight, then were washed twice with PBS. Prior to lysis, dishes were washed with ice-cold PBS. Cells were then lysed in 50mM Tris pH 7.4, 150 mM NaCl, 1mM EDTA, 1% Triton X-100 plus protease and phosphatase inhibitor cocktails (Complete Mini, EDTA-free; PhosSTOP; Roche Diagnostics). Insoluble material was cleared by centrifugation for 10 min at 20,000xg. Immunoprecipitations were performed on the resulting lysates. For anti-HA immuno-precipitations, anti-HA Affinity Matrix (Roche Diagnostics) was used. For anti-FLAG immunoprecipitations, anti-FLAG M2 affinity gel was used (Sigma-Aldrich). Lysates were added to resin and incubated for 1 hour at 4°C with rotation. Afterwards, resin was washed five times with lysis buffer, then eluted with 2x Laemmli buffer at 50°C for three minutes. Laemmli buffer was removed from resin and 6.25% 2-mercaptoethanol was added. Immunoblotting was performed with 4–15% gradient Mini-PROTEAN TGX precast polyacrylamide gels and nitrocellulose membranes (Bio-Rad). Blots were blocked with 5% non-fat dry milk (AmericanBIO), and antibodies were incubated with 5% non-fat dry milk or bovine serum albumin (AmericanBIO) in TBS with 0.1% Tween 20. Chemiluminescence detection of horseradish peroxidase signals from secondary antibodies was performed on a Versa-Doc imaging station (Bio-Rad). Antibodies used in this study are listed in Supplemental Table 3.

### Mass spectrometry

HeLa cells expressing Lyso-C9orf72-GFP and SMCR8-tdTomato or Lyso-GFP and SMCR8-tdTomato were lysed as described above. Lysates were incubated with GFP-Trap resin (ChromoTek) for 2 h at 4°C. After washing in lysis buffer, bound proteins were eluted with 2xLaemmli buffer and heated at 37°C. Samples were then run 10 mm into an SDS-PAGE gel, silver stained and excised from the gel.

Silver-stained gel bands were treated with 5% acetic acid for 10 minutes with rocking. The acid was removed and the bands were covered with freshly prepared destaining solution (1:1 ratio of stock solutions of 30 mM potassium ferricyanide in water and 100 mM sodium thiosulfate in water) until the brownish color disappeared. The bands were then rinsed three times with 0.5 mL water for 5 minutes to remove the acid and chemical reducing agents. The gel bands were cut into small pieces and washed for 30 minutes on a tilt-table with 450 μL 50% acetonitrile/100 mM NH4HCO3 (ammonium bicarbonate) followed by a 30-minute wash with 50% acetonitrile/12.5 mM NH4HCO3. The gel bands were shrunk by the brief addition then removal of acetonitrile, then dried by speed vacuum. Each sample was resuspended in 100 μL of 25 mM NH4HCO3 containing 0.5 μg of digestion grade trypsin (Promega, V5111) and incubated at 37 °C for 16 hours. Supernatants containing tryptic peptides were transferred to new Eppendorf tubes and the gel bands were extracted with 300 μL of 80% acetonitrile/0.1% trifluoroacetic acid for 15 min. Supernatants were combined and dried by speed vacuum. Peptides were dissolved in 24 μL MS loading buffer (2% aceotonitrile, 0.2% trifluoroacetic acid), with 5 μL injected for LC-MS/MS analysis.

LC-MS/MS analysis was performed on a Thermo Scientific Q Exactive Plus equipped with a Waters nanoAcquity UPLC system utilizing a binary solvent system (A: 100% water, 0.1% formic acid; B: 100% acetonitrile, 0.1% formic acid). Trapping was performed at 5 μL min^−1^, 97% Buffer A for 3 min using a Waters Symmetry C18 180 μm × 20 mm trap column. Peptides were separated using an ACQUITY UPLC PST (BEH) C18 nanoACQUITY Column 1.7 μm, 75 μm × 250 mm (37 °C) and eluted at 300 nl/min with the following gradient: 3% buffer B at initial conditions; 5% B at 1 minute; 35% B at 90 minutes; 50% B at 105 minutes; 90% B at 110 minutes; 90% B at 115 min; return to initial conditions at 116 minutes. MS was acquired in profile mode over the 300-1,700 m/z range using 1 microscan, 70,000 resolution, AGC target of 3E6, and a maximum injection time of 45 ms. Data dependent MS/MS were acquired in centroid mode on the top 20 precursors per MS scan using 1 microscan, 17,500 resolution, AGC target of 1E5, maximum injection time of 100 ms, and an isolation window of 1.7 m/z. Precursors were fragmented by HCD activation with a collision energy of 28%. MS/MS were collected on species with an intensity threshold of 2E4, charge states 2-6, and peptide match preferred. Dynamic exclusion was set to 20 seconds.

Tandem mass spectra were extracted by Proteome Discoverer software (version 2.2.0.388, Thermo Scientific) and searched in-house using the Mascot algorithm (version 2.6.0, Matrix Science). The data were searched against the SwissProt database (version 2017 09) with taxonomy restricted to Homo sapiens (20,238 sequences). Search parameters included trypsin digestion with up to 2 missed cleavages, peptide mass tolerance of 10 ppm, MS/MS fragment tolerance of 0.02 Da, and methionine oxidation as a variable modification. Normal and decoy database searches were run, with the confidence level was set to 95% (p<0.05). Scaffold (version Scaffold 4.8.9, Proteome Software) was used to validate and compare MS/MS based peptide and protein identifications.

### Immunofluorescence and microscopy

A Nikon Inverted Eclipse TI-E Microscope equipped with 60x CFI PlanApo VC, NA 1.4, oil immersion and 40x CFI Plan Apo, NA 1.0, oil immersion objectives, a spinning disk confocal scan head (CSU-X1, Yokogawa) and Volocity (PerkinElmer) software was used for spinning-disk confocal microscopy. Images were acquired at room temperature (22°C). CCells were grown on 12 mm No. 1.5 coverslips (Carolina Biological Supply, Burlington NC) coated with Poly-D lysine (Sigma-Aldrich) and fibronectin (EMD-Millipore). Cells were fixed by adding 1 volume of 8% paraformaldehyde (PFA) in 0.1 M sodium phosphate to the culture media in a dropwise manner in order to achieve a final concentration of 4% PFA. Cells were fixed at room temperature for 30 minutes, then washed with phosphate buffered saline (PBS). Samples were permeabilized by immersing coverslips in ice-cold methanol for three seconds, followed by PBS rinses. Samples were then blocked in 5% normal donkey serum (Jackson ImmunoResearch)/PBS for one hour at room temperature. All subsequent antibody incubations were performed in this buffer. Antibodies used in this study are listed in Supplemental Table 3. TThe protocol for detecting 2xHA-tagged endogenously expressed protein was performed as described previously (Petit et al., 2013). The percentage of starved cells with C9orf72 puncta colocalized with LAMP1 in Figure 2 was made by visual inspection of images from three independent experiments with >140 cells per cell line analyzed. Images were processed with ImageJ (National Institutes of Health).

### CRISPR/Cas9 genome editing

Guide RNA targeting exon 2 of PQLC2 and exon 3 of *CTNS* (the gene encoding cystinosin) were selected from predesigned gRNA sequences (Wang et al., 2014). Cloning of guide RNA–encoding DNA oligonucleotides (Integrated DNA Technologies) into Bbs1-digested pX459 vector was done as described previously (Amick et al., 2018; Ran et al., 2013). Oligonucleotide sequences for generating the gRNA used to target *PQLC2* and *CTNS* are listed in Supplemental Table 2. Sequences of the guide RNA plasmids were confirmed by sequencing. Nprl3 gRNA sequences are described elsewhere (Meng and Ferguson, 2018). 0.4 μg plasmid DNA was transfected with FuGENE 6 into 250,000 cells in a 6-well dish. The next day, transfected cells were selected with 1.25 μg mL^−1^ puromycin for 3 days. Surviving cells were subsequently plated at clonal density. Following the selection and expansion of colonies, PQLC2 knockouts were identified by sequencing of PCR-amplified genomic DNA. To sequence genomic DNA, it was extracted (QuickExtract DNA extraction solution, Epicentre Biotechnologies), the region of interest was amplified by PCR (primers described in Supplemental Table 4), cloned into the pCR-Blunt TOPO vector (Zero Blunt TOPO PCR cloning kit, ThermoFisher Scientific), and transformed into TOP10-competent E. coli cells. Plasmid DNA was then isolated from colonies and sequenced to define the genotype of the locus of interest.

The method used for CRISPR/Cas9 genome editing to insert epitope tags at endogenous loci was described previously (Amick et al., 2018). The single strand DNA oligonucleotide (ssODN) homology-directed repair donor template was designed with asymmetric homology arms on the PAM proximal and distal sides of the cut site in order to enhance the efficiency of tag insertion (Richardson et al., 2016). The ssODN repair template and protospacer sequences (Integrated DNA Technologies) used to insert the 2xHA epitope tag into the PQLC2 gene are listed in Supplemental Table 5. Clonal cell populations were isolated and screened for HA signal by immunoblotting and immunofluorescence. To confirm further the correct in-frame insertion of the 2xHA tag in these cells, genomic DNA surrounding the site of tag insertion was PCR amplified and sequenced as described above. Primer sequences used for this purpose are listed in Supplemental Table 4.

### Magnetic isolation of lysosomes

Magnetic isolation of lysosomes from cells that were endocytically loaded with colloidal iron dextran nanoparticles was performed as described previously (Amick et al., 2018). Lysosomes were purified from the following cell lines under starved conditions: HEK293FT cells expressing 2xHA-C9orf72 from the endogenous locus, PQLC2 knockout in the background of the aforementioned cell line, PQLC2 knockout cells stably expressing wild type PQLC2-FLAG and cells stably expressing P55L/P201L PQLC2-FLAG. Lentiviral transduction was used to achieve stable transgenic expression of PQLC2-FLAG and P55L/P201L mutant. Generation of plasmids for this purpose is described above. Lentivirus packaging and infection was performed as described previously (Amick et al., 2018). Magnetic isolation of lysosomes from cells that were endocytically loaded with colloidal iron dextran nanoparticles was performed as described previously (Amick et al., 2018; Tharkeshwar et al., 2017). Lysosomes were purified from the following cell lines under starved conditions: HEK293FT cells expressing 2xHA-C9orf72 from the endogenous locus, PQLC2 knockout in the background of the aforementioned cell line, PQLC2 knockout cells stably expressing wild type PQLC2-FLAG and cells stably expressing P55L/P201L PQLC2-FLAG. Lentiviral transduction was used to achieve stable transgenic expression of PQLC2-FLAG and P55L/P201L mutant. Generation of plasmids for this purpose is described above. Lentivirus packaging and infection was performed as described previously (Amick et al., 2018).

### Amino acid starvations and re-feeding

For cells analyzed in fed conditions, cells were incubated with fresh normal growth media (DMEM) as described above for two hours prior to analysis. For starvation experiments, dishes were washed with PBS, then incubated in RPMI medium without amino acids, without serum, with glucose (United States Biological). Where indicated, dialyzed FBS was added to this media (Gibco by Life Technologies). Also where indicated, minimum essential media (MEM) amino acids solution (Gibco) was added to this media. For experiments in Fig. 4A, cells were incubated in media with or without these components for two hours prior to lysis. The MEM amino acids solution at 1x contains L-arginine (600μM), L-cystine (100 μM), L-histidine (200 μM), L-isoleucine (400 μM), L-leucine (400 μM), L-lysine (396 μM), L-methionine (101 μM), L-phenylalanine (200 μM), L-threonine (400 μM), L-tryptophan (50 μM), L-tyrosine (199 μM) and L-valine (400 μM).) Mixtures of individual amino acids were prepared to yield the “RKH” and “Other” amino acids at their respective concentrations. For experiments in Fig. 4F, cells were incubated in starvation media for two hours, followed by the incubation in media containing the indicated groups of amino acids for one hour. . For experiments in Fig. 4D, cells were incubated in media lacking the indicated amino acids and containing the others at their respective concentrations.

### Cystinosin depletion and cysteamine treatment

Guide RNAs used to target the CTNS gene and the method of generating a cystinosin-depleted clonal cell line in the background of PQLC2-2xHA cells is described in the CRISPR/Cas9 genome editing section and Supplemental Table 2. These cells (and the parental PQLC2-2xHA cell line) were starved as described above, then cysteamine (Sigma-Aldrich) was added to starvation media at 1 mM final concentration for 1 hour prior to lysis. Immunoprecipitations and immunoblotting were performed as described in previous sections.

### Statistical analysis

Data were analyzed using Prism (GraphPad software) and specific statistical tests are specified in the figure legends. All error bars represent SEM. Data distribution was assumed to be normal, but this was not formally tested.

## Supplemental Material

Fig. S1 shows the localization of FLAG-tagged PQLC2 and PAT1. Fig S2 shows the localization of C9orf72 in GATOR1-deficient cells. Fig S3 shows the localization of PQLC2 mutants. Fig. S4 shows the 2xHA epitope tagging of the endogenous PQLC2. Fig. S5 shows the localization of PQLC2 under different nutrient conditions. Tables S1 and S2 contain DNA sequences used for generating the plasmids used in this study. Table S3 contains the antibodies used in this study. Table S4 contains the primers used for PCR amplification of genomic DNA. Table S5 contains the sequences used for the PQLC2 CRISPR knock-in.

---

**Supplemental Figure 1.**
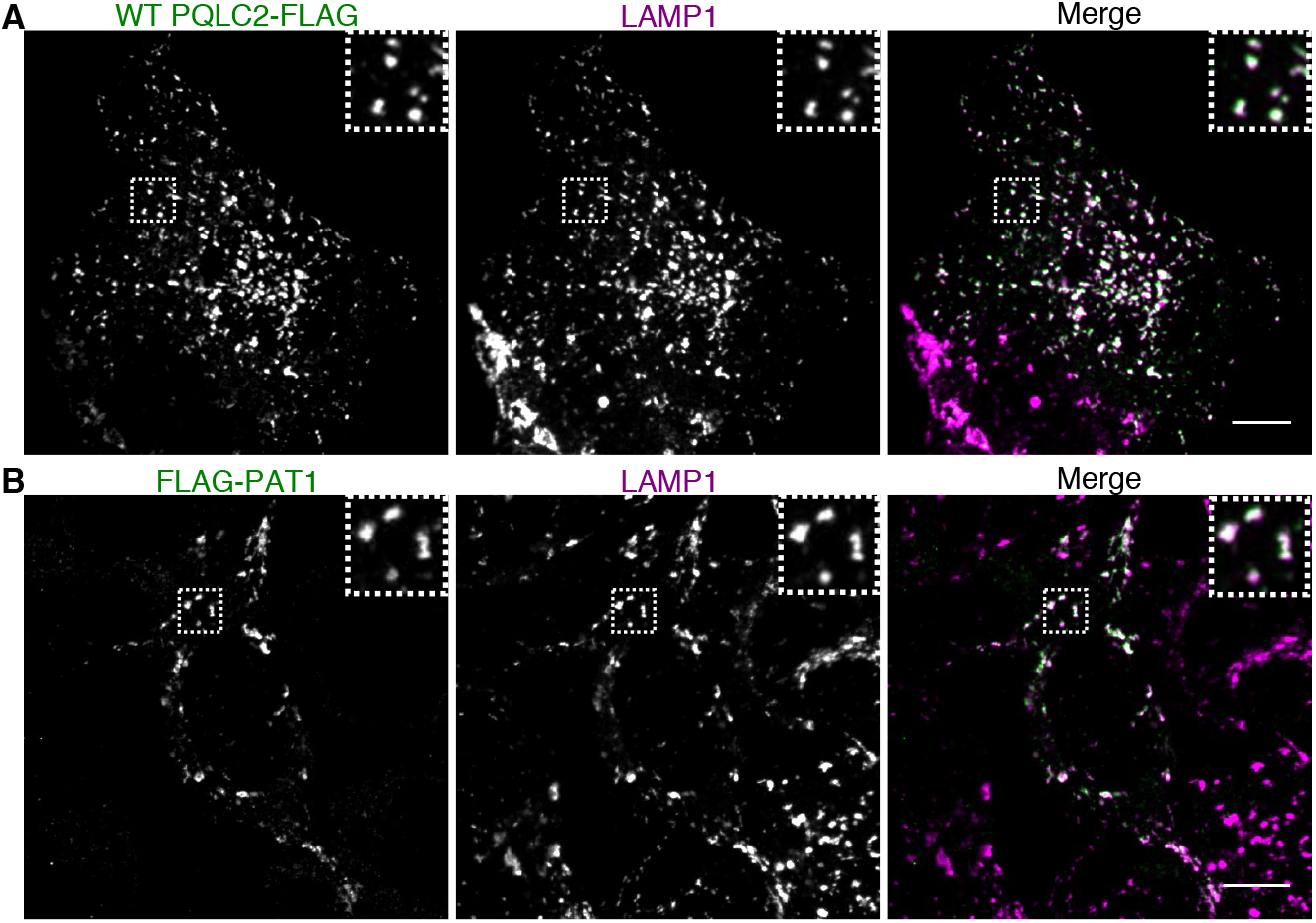
PQLC2-FLAG and FLAG-PAT1 localize to lysosomes. Immunofluorescence images of FLAG and LAMP1 in HEK293FT cells transiently transfected with FLAG-tagged PQLC2 **(A)** or PAT1 **(B)**. Scale bars: 10 μm.

**Supplemental Figure 2.**
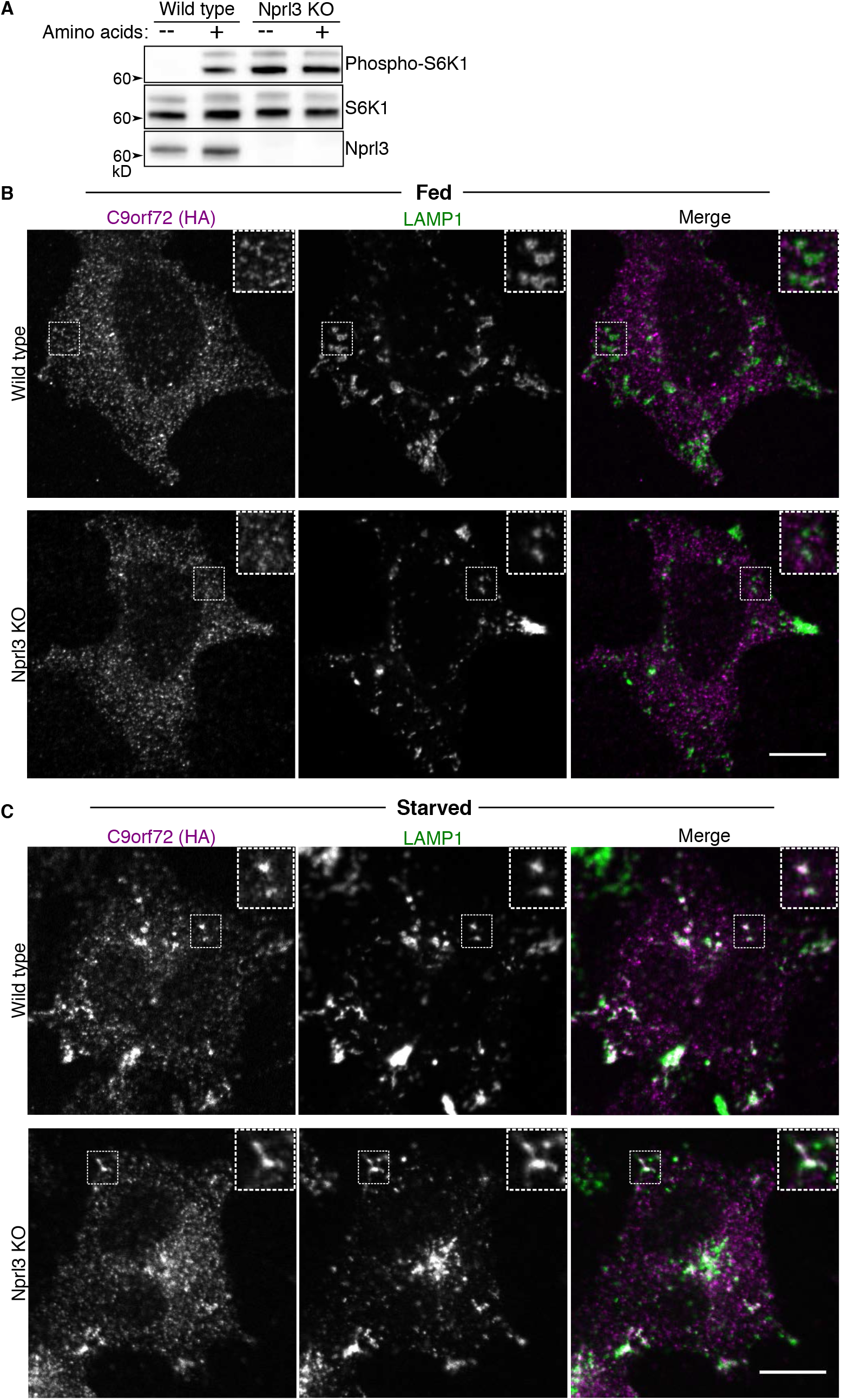
C9orf72 recruitment to lysosomes in independent of GATOR1-associated nutrient sensing. **(A)** Immunoblot analysis of Nprl3, S6 kinase and phospho S6K levels during starvation (2 h) and amino acid refeeding (15 min) in wild type and Nprl3 KO cells. **(B)** Immunofluorescence images of C9orf72 localization under normal, fed growth conditions for the indicated genotypes. **(C)** Immunofluorescence images of C9orf72 localization under starved (2 h) conditions for the indicated genotypes. Localization of C9orf72 to lysosomes is observed by its colocalization with LAMP1. Scale bars: 10 μm.

**Supplemental Figure 3.**
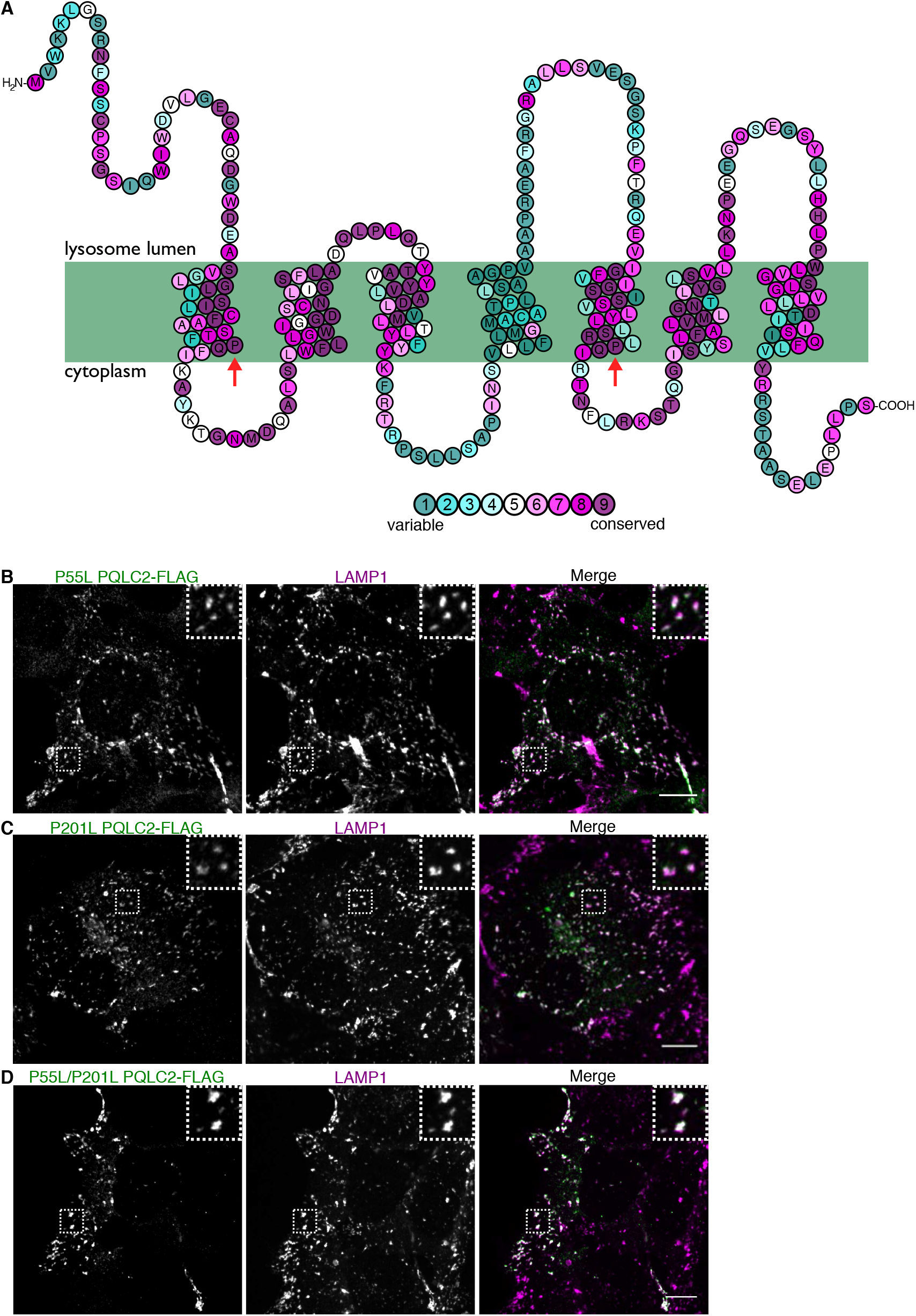
PQLC2-FLAG PQ loop mutants localize to lysosomes. **(A)** Visualization of PQLC2 topology generated by Protter overlaid with the conservation of amino acids at each position from ConSurf (Ashkenazy et al., 2016). The position of conserved prolines in the PQ loops are indicated by arrows. **(B-D)** Immunofluorescence images of FLAG and LAMP1 in HEK293FT cells transiently transfected with FLAG-tagged PQLC2 mutants: P55L **(B)**, P201L **(C)** and P55L/P201L **(D)**. Scale bars: 10 μm.

**Supplemental Figure 4.**
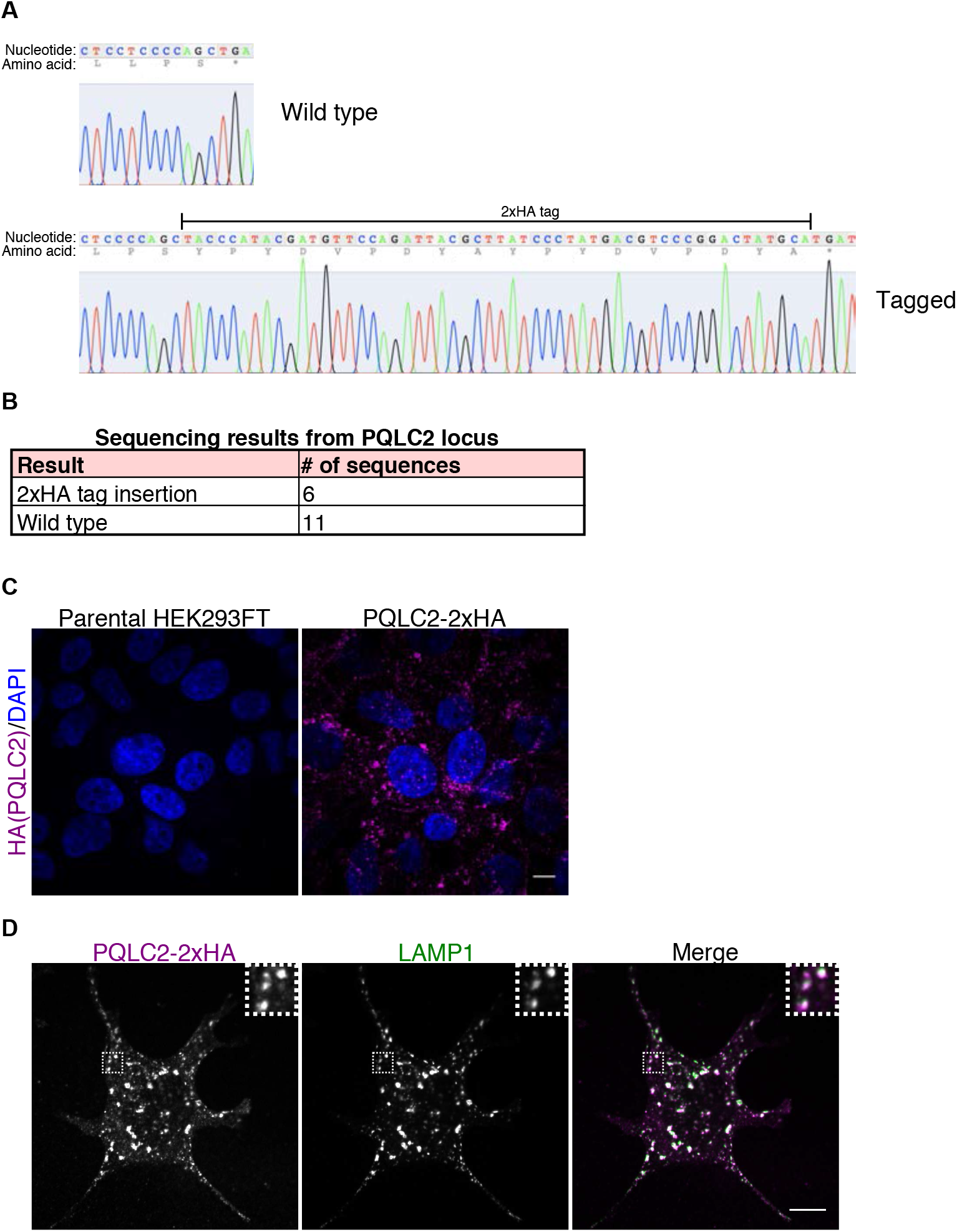
Epitope tagging of the endogenous PQLC2 protein. **(A)** Sequencing traces from an HEK293FT cell line that has a 2xHA epitope tag inserted at the endogenous locus. Example traces from unaffected and edited alleles are shown. The 2xHA tag is at the C-terminus of PQLC2. **(B)** Summary of sequencing results from this cell line. Results are consistent with one of three copies of PQLC2 being 2xHA-tagged at the C-terminus, while two of three are unaffected. **(C)** The specificity of the anti-HA immunofluorescence signal in PQLC2-2xHA cells is supported by the absence of this signal in parental, non gene-edited cells. **(D)** Immunofluorescence images showing the localization of PQLC2-2xHA to lysosomes (endogenous PQLC2-2xHA and LAMP1). Scale bars: 10 μm.

**Supplemental Figure 5.**
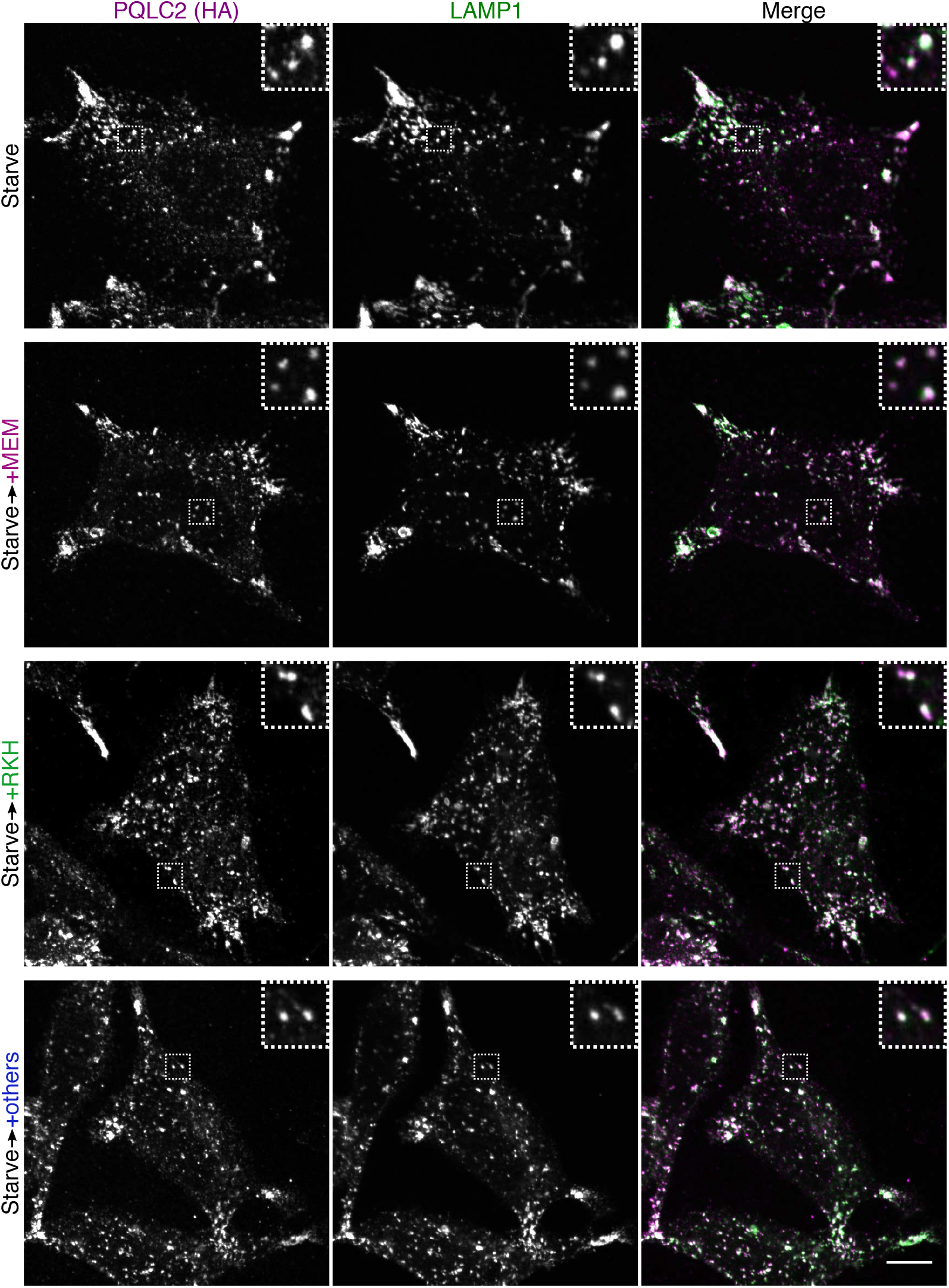
Amino acid availability does not noticeably alter PQLC2 localization. Immunofluorescence images of PQLC2-2xHA and LAMP1 in cells expressing PQLC2-2xHA from the endogenous locus. Cells were starved, then incubated in media containing the indicated amino acids (as in Figs. 4F and 5). Scale bar: 10 μm.

**Supplemental Table 1.**
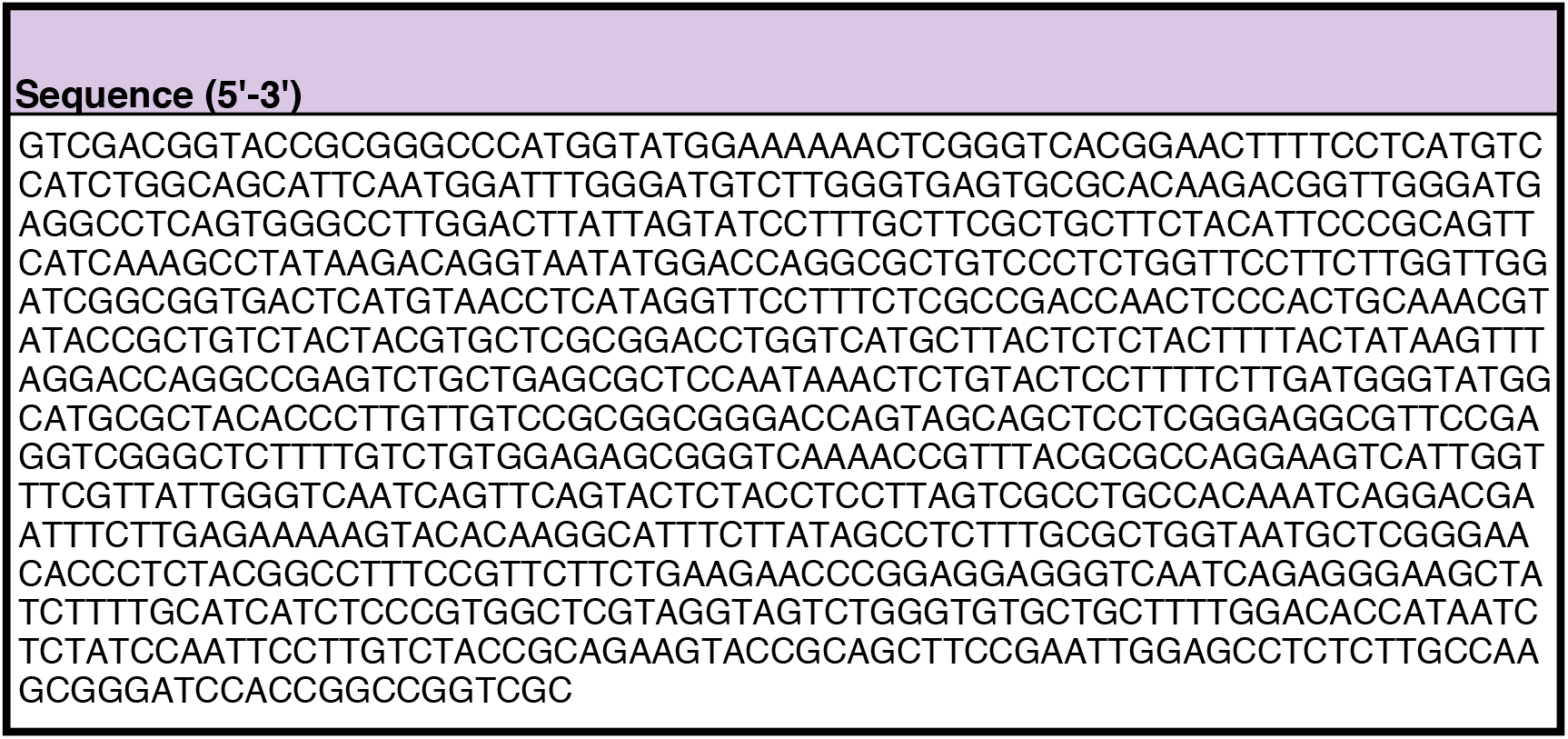
PQLC2 synthetic DNA fragment sequence for cloning into SmaI-digested vector.

**Supplemental Table 2.**
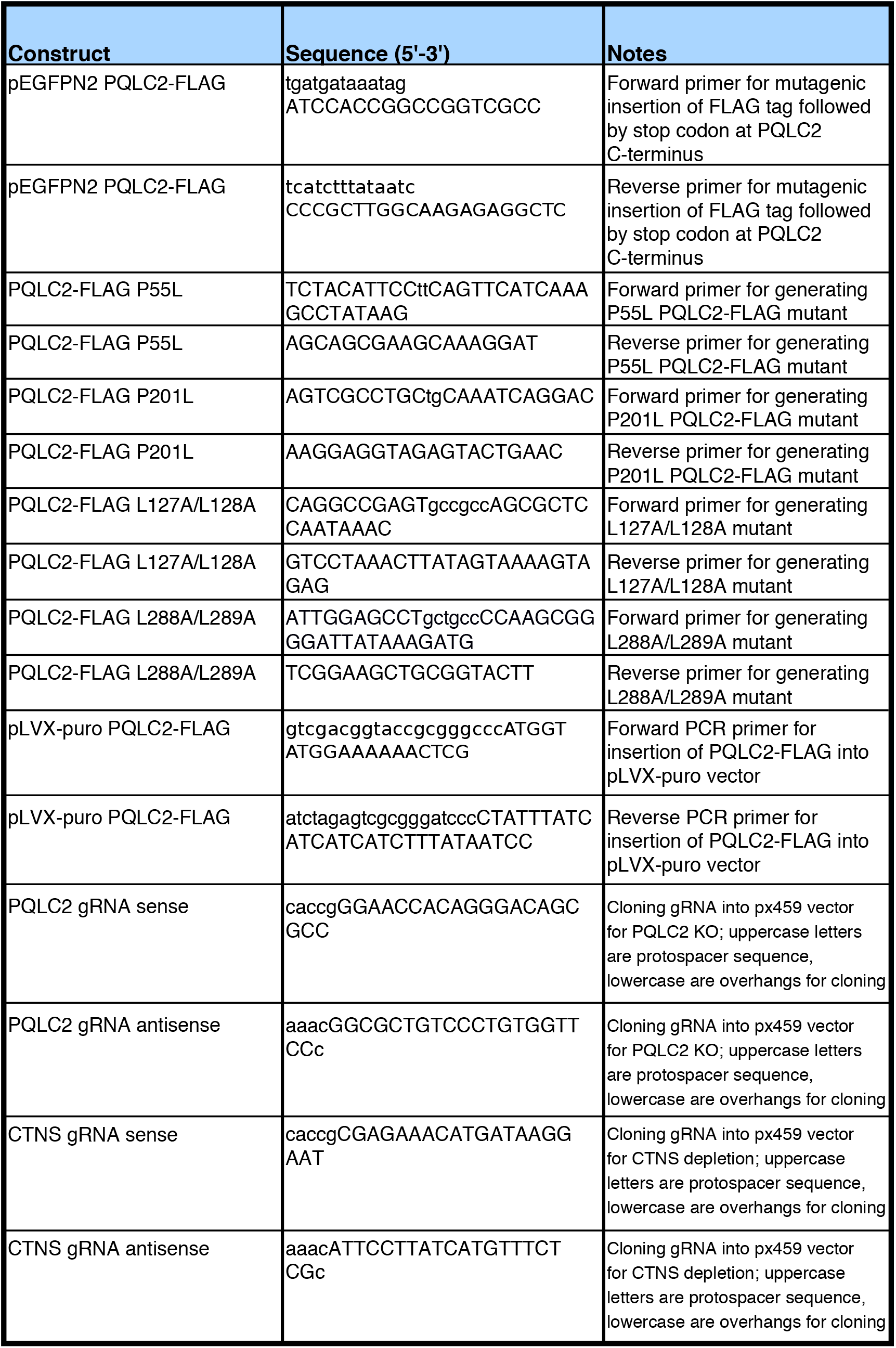
PCR primers and other oligo sequences.

**Supplemental Table 3.** Summary of antibodies used.

| **Antibody** | **Source** | **Catalog/clone number** |
| --- | --- | --- |
| C9orf72 | Proteintech | 22637-1-AP |
| Cystinosin | Abnova | H00001497-M09 |
| Donkey anti-Mouse,<br>Alexa Fluor 594 | Thermo Fisher Scientific | A-21203 |
| Donkey anti-Goat,<br>Alexa Fluor 594 | Thermo Fisher Scientific | A-11058 |
| Donkey anti-Rabbit,<br>Alexa Fluor 488 | Thermo Fisher Scientific | A-21206 |
| FLAG (DYKDDDDK) | Sigma-Aldrich | F7425 |
| FLAG (DYKDDDDK) | Cell Signaling Technology | 2368S |
| FLCN | Cell Signaling Technology | 3697 |
| Goat Anti-Mouse IgG | Jackson ImmunoResearch | 115-005-146 |
| HA | Cell Signaling Technology | 2367 |
| HA-Peroxidase (3F10) | Roche | 12 013 819 001 |
| LAMP1 | Cell Signaling Technology | 9091 |
| LAMP1 | University of Iowa DSHB | H4A3 |
| Nprl3 | Sigma-Aldrich | HPA011741 |
| phospho-p70 S6<br>Kinase (Thr389) | Cell Signaling Technology | 9234 |
| p70 S6 Kinase | Cell Signaling Technology | 9202 |
| SMCR8 | Bethyl Laboratories | A304-694A |
| WDR41 | Novus Biologicals | NBP1-83812 |

**Supplemental Table 4.**
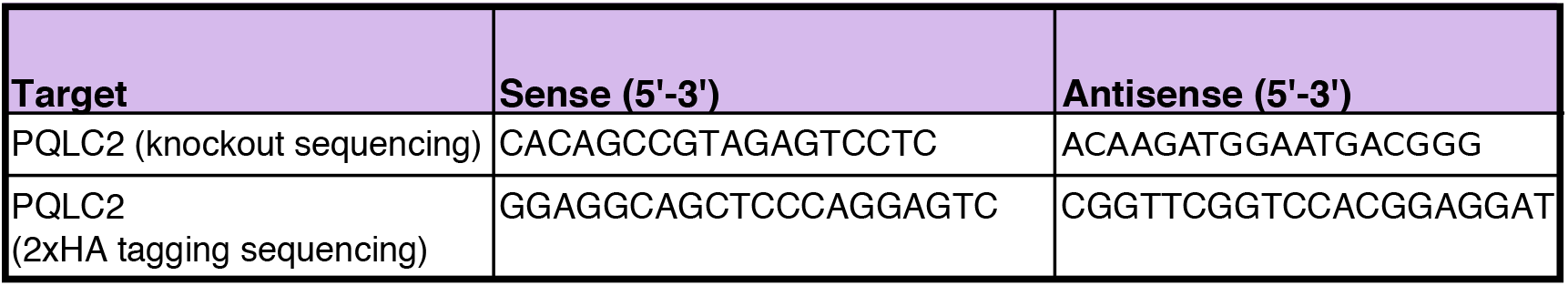
PCR primers for genomic DNA PCR.

**Supplemental Table 5.**
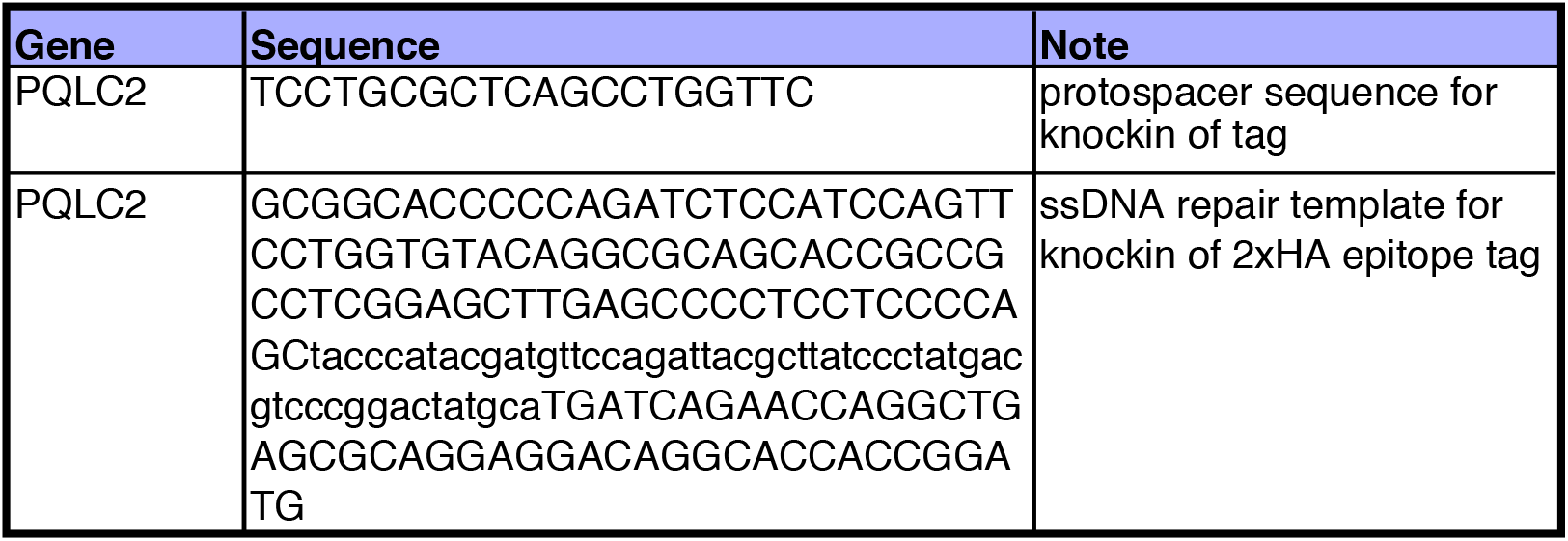
Sequences used for CRISPR knockin of 2xHA epitope tag into PQLC2 locus.

